# Patchy and widespread distribution of bacterial translation arrest peptides associated with the protein localization machinery

**DOI:** 10.1101/2023.09.02.556018

**Authors:** Keigo Fujiwara, Naoko Tsuji, Mayu Yoshida, Hiraku Takada, Shinobu Chiba

## Abstract

Regulatory arrest peptides exert cellular functions via mechanisms involving regulated translational arrest. Monitoring substrates, a class of arrest peptides, feedback-regulate the expression of the Sec or YidC protein localization machinery. Previously, only a limited number of monitoring substrates were identified. In this study, we performed a bacterial domain-wide search, followed by *in vivo* and *in vitro* analyses, leading to a comprehensive identification of many novel Sec/YidC-related arrest peptides that showed patchy, but widespread, phylogenetic distribution throughout the bacterial domain. Identification of five novel arrest-inducing sequences suggests that bacteria have evolved various arrest-inducing mechanisms. We also identified many arrest peptides that share an R-A-P-P like sequence, suggesting that this sequence could serve as a common evolutionary seed that could overcome the species-specific structures of ribosomes, to evolve arrest peptides. Our comprehensive phylogenetic study revealed that arrest peptide is a prevalent mechanism for the gene regulation of the protein localization machinery.

## Introduction

Regulatory nascent chains exert their cellular functions while they are still nascent polypeptides^1–3^. They induce programmed ribosomal stalling by interacting with ribosomal residues located near the peptidyl transferase center (PTC), the mid-tunnel region within the nascent polypeptide exit tunnel (NPET), and, occasionally, on the ribosomal surface^4–8^. Thus, they are also called ribosome arrest peptides (RAPs) or simply arrest peptides^1^. Translation arrest generally occurs under a specific intracellular condition, thus allowing arrest peptides to respond to changes in the intracellular environment to serve as sensors of the feedback gene regulation.

A class of bacterial arrest peptides, such as SecM, MifM, and VemP, is involved in the feedback regulation of genes encoding components of the protein localization machinery^9–11^. *Escherichia coli* SecM and *Vibrio alginolyticus* VemP monitor the Sec protein secretion pathway, in which SecA and SecDF facilitate protein translocation in ATP– and proton-motive force-depending manners, respectively^12–15^. *Bacillus subtilis* MifM monitors the YidC membrane protein insertion pathway, in which YidC serves as an “insertase” for a class of membrane proteins^16–20^.

The *secM* gene encodes a protein with the N-terminal Sec-dependent signal sequence^21^ and the C-terminal arrest sequence (F_150_xxxxWIxxxxGIRAGP_166_)^22^, and is co-transcribed with its downstream *secA* gene (Fig. 1a)^23^. A stem-loop structure sequesters the Shine–Dalgarno (SD) sequence of *secA*^24^. The stalled ribosome on *secM* interferes with the stem-loop structure, thus allowing SecA synthesis^22,25^. Engagement of the SecM nascent chain with the active Sec translocation machinery leads to the arrest cancellation^26^. Thus, a malfunction of the Sec machinery results in a prolonged arrest of SecM, leading to *secA* induction^25^. MifM and VemP feedback-regulate the downstream *yidC2* (*yqjG*) and *secD2/F2* genes, respectively, in a similar fashion^10,11^. As SecM, MifM, and VemP are substrates of the protein-translocation pathway that they monitor, they are also called “monitoring substrates”^27^.

**Figure 1:**
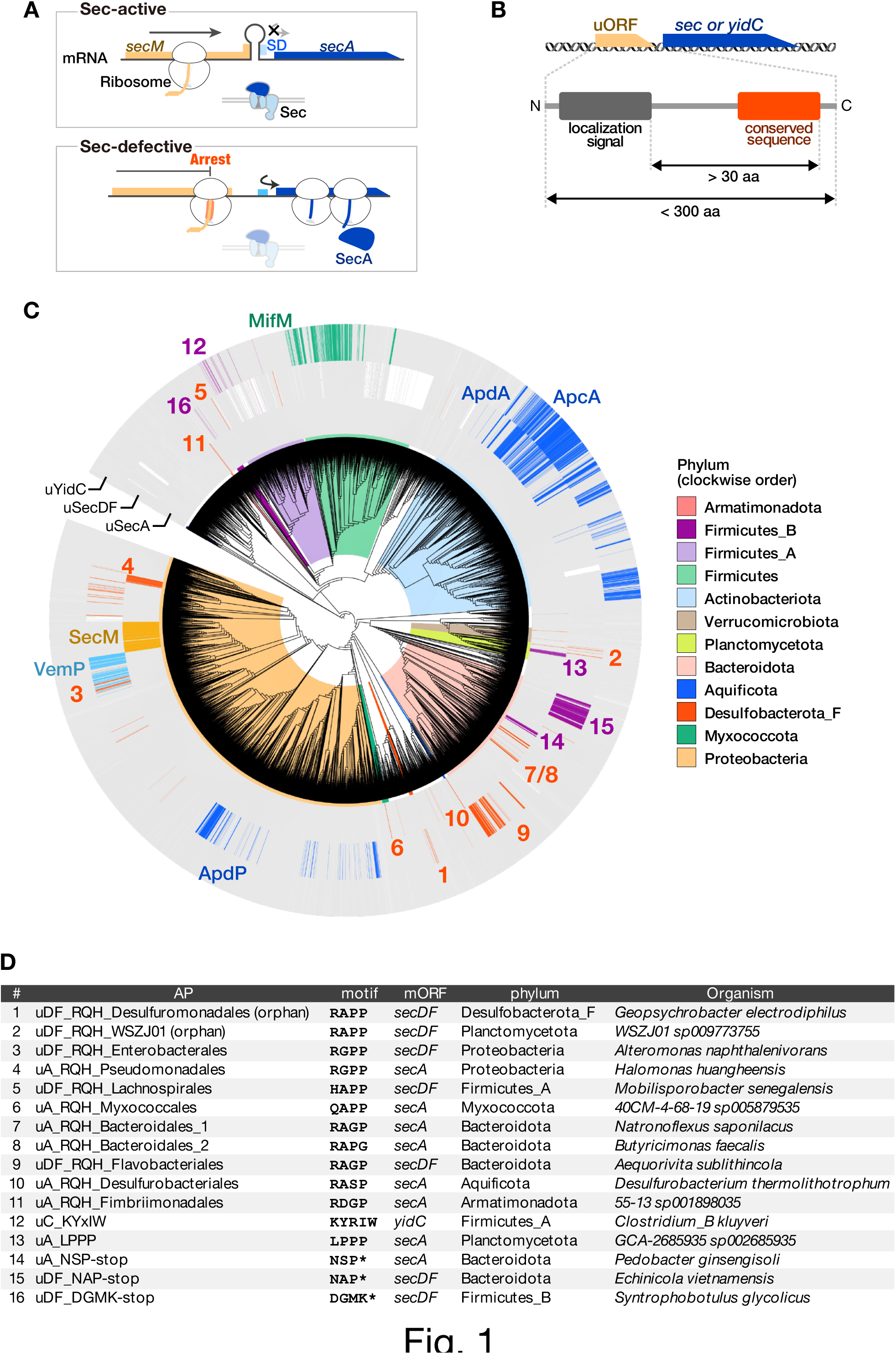
Patchy and widespread phylogenetic distribution of candidate monitoring substrates across the bacterial tree of life. **a**, The translation arrest of SecM induces the *secA* gene by disrupting the stem-loop structure that otherwise sequesters SD. **b,** Searching criteria for novel monitoring substrates. **c,** Phylogenetic distribution of the candidate monitoring substrates. The gray strips around the bacterial phylogenetic tree indicate the genomes encoding SecA, SecD/F, or YidC homologs. Genomes with genes encoding putative arrest peptides located upstream of the *secA*, *secDF*, or *yidC* gene are indicated by chromatic strips, with the names or numbers corresponding to those reported in the list shown in (d). The colors behind the tree indicate the bacterial phylum listed on the right. **d,** General information of the representative candidate arrest peptides used in *in vivo* and *in vitro* experiments. The numbers correspond to those in (c).

The lack of sequence similarity among the arrest sequences has hampered the identification of novel arrest peptides based on conventional approaches. To overcome this obstacle, we recently established an *in silico* screening system to find open reading frames (ORFs) that possibly encode monitoring substrates based on features shared by known monitoring substrates^28^. Our previous search across 449 bacterial genomes identified three arrest peptides, i.e., ApcA and ApdA from actinobacteria and ApdP from Alphaproteobacteria^28^. *apcA* was located upstream of *yidC2*, whereas *apdA* and *apdP* were located upstream of *secDF2*. Notably, crucial residues located near arrest sites exhibited a sequence similarity: ApdA and ApdP harbored the R-A-P-P sequence, whereas ApcA harbored the R-A-P-G sequence, which was also reminiscent of the R-A-G-P sequence corresponding to the C-terminal part of the arrest motif of SecM from *E. coli*^22,28^.

Since we found that the screening described above could potentially be a groundbreaking approach to the identification of novel arrest peptides, we envisioned that we could comprehensively identify arrest peptides and depict a domain-wide phylogenetic view by applying it to a larger and comprehensive set of genome databases.

In this study, we conducted a systematic search for monitoring substrates to understand bacterial domain-wide nature of the arrest sequence. We utilized over 30,000 representative bacterial genomes of Genome Taxonomy Database (GTDB)^29^, to guarantee the comprehensiveness. Strikingly, our current screening led to the identification of a large number of homology groups encoded upstream of the *sec*A, *secDF*, and *yidC* genes. Interestingly, many of the ORFs identified bore RAPP-like sequences, such as RAPP, RGPP, RAGP, and RAGP. Furthermore, we identified several ORFs encoding distinct conserved sequences near the C-termini. Our subsequent *in vivo* and *in vitro* analyses provided evidence that they were able to efficiently stall the ribosomes of either *E. coli* or *B. subtilis*, or both. We also demonstrated that one of the arrest peptides identified here induced the downstream *secDF* gene in a translation arrest-dependent manner. These data suggest that a wide variety of bacteria have evolved arrest peptides encoded by uORFs of the *sec* or *yidC* genes to regulate these genes.

## Results

### Bioinformatics search for novel arrest peptides

To address the extent to which the arrest peptide-mediated regulation of genes involved in the protein localization machinery is universal in bacteria, we first carried out an in-depth *in silico* screening of ORFs encoding novel translation arrest peptides using over 30,000 representative bacterial genomes (Table S1) from the GTDB^29^. We used the following search criteria, which we employed previously to identify ORFs encoding arrest peptides (Fig. 1b)^28^: (i) a uORF of the *sec* or *yidC* gene encoding a small protein with no annotated function; (ii) a uORF encoding an N-terminal secretion signal or a transmembrane (TM) sequence; (iii) a uORF encoding a C-terminal sequence conserved among its homologs; and (iv) a uORF encoding a spacer region between the localization signal and the C-terminal end of the conserved sequence with a size greater than 30 amino acid residues to ensure the exposure of the N-terminal localization signal outside of the ribosome when arrested. We extracted short ORFs of unknown function located upstream of the *secA*, *secDF*, and *yidC* genes and classified them into groups via clustering based on their amino acid sequences using the MMseqs2 function^30^. Subsequently, we collected candidate clusters that met the criteria described above (see Materials and Methods).

This *in silico* screening allowed us to identify several homologs of known arrest peptides, as well as many candidate ORFs that possibly encode novel arrest peptides (Fig. 1c; Supplementary Fig. 1-12; Supplementary Tables 2–4). Strikingly, a substantial number of uORFs that met our criteria encompassed C-terminal motifs that were similar to that of SecM (R-A-G-P), ApcA (R-A-P-G), or ApdA/ApdP (R-A-P-P). We also identified uORFs that encoded similar C-terminal motifs, such as R-G-P-P, H-A-P-P, and Q-A-P-P. In this study, we refer to these uORFs (i.e., those sharing the RAPP-like sequence that were not homologs of SecM, ApcA, ApdA, or ApdP) as the RQH family, as per the first residues of the consensus motifs. The RQH family members were widely, but also patchily, distributed among nine independent phyla (Supplementary Fig. 1). This contrasted with the ubiquitously distributed *secA*, *secDF*, and *yidC* genes (Fig. 1c, gray bars). The lack of an overall sequence similarity among the RQH family members hampered their rational categorization into homology groups using conventional means. Therefore, we divided them into 19 groups based on the bacterial order in which they were identified, and provisionally named each of them based on the downstream gene, motif code (RQH, in this case), and bacterial order name using the following rule: “[uA/uDF/uC, which indicate a uORF of *secA*, *secDF*, and *yidC*, respectively]_[representative residues]_[bacterial order].” For instance, members of the RQH family identified upstream of *secA* in a subset of the order Pseudomonadales are referred to as uA_RQH_Pseudomonadales (Fig. 1c and d). Among the 19 groups of the RQH family, eight groups were detected upstream of *secA*, whereas the remaining 11 groups were observed upstream of *secDF* (Supplementary Fig. 1). In addition, we identified several uORFs that harbored the RAPP/RGPP motif but were apparently not conserved among their respective bacterial orders, possibly because of their too-narrow phylogenetic distributions. These orphan uORFs with RAPP/RGPP motifs were detected in nine phyla (Supplementary Fig. 1). Thus, for further *in vivo* and *in vitro* analyses, we selected nine representative uORFs from eight major RQH family groups, as well as two orphan uORFs (Fig. 1c and d).

Furthermore, we identified several clusters of uORFs that encoded a unique consensus motif apparently unrelated to that of any known arrest peptides. These uORFs were also provisionally named according to their downstream gene and conserved motif ([uA/uDF/uC]_[conserved residues]), as exemplified by uA_LPPP, which shares the L-P-P-P motif near the C-terminus (Fig. 1c, d and Supplementary Fig. 7). The members of the uC_KYxIW cluster detected in the Clostridia class of Firmicutes_A typically shared a unique K-Y-x-I-W sequence (x = any residue) (Supplementary Fig. 3). Several uORF clusters that shared a proline at the C-terminal end were identified exclusively in Bacteroidota (Fig. 1c, d and Supplementary Fig. 6). Members of the uA_NSP-stop and uDF_NAP-stop clusters encoded peptides encompassing N-S-P and N-A-P motifs, respectively, at their C-terminal ends. (Fig. 1c, d and Supplementary Fig. 6). Finally, uDF_DGMK-stop was observed in subsets of Firmicutes_A and Firmicutes_B (Fig. 1c, d and Supplementary Fig. 3).

### In vitro translation arrest of the RQH family members

Next, we addressed whether these candidate arrest peptides stall the ribosome using the PURE system, a coupled *in vitro* transcription–translation system with all purified components derived from *E. coli* (*Ec* PURE)^31^, as well as the *Bs* hybrid PURE system (*Bs* PURE)^32^, in which only the ribosome of *Ec* PURE is replaced by the *B. subtilis* ribosome. For the *in vitro* translation assay, a gene segment encoding the C-terminal soluble region of a candidate arrest peptide (*ap*) was sandwich-fused between the *gfp* and *lacZα* genes (Fig. 2a). Ribosome stalling would result in the accumulation of the N-terminal GFP-AP fragment with a covalently bonded tRNA at the C-terminal end, which would migrate even slower than the full-length GFP-AP-LacZα fragment on SDS–PAGE, unless the tRNA moiety is removed by RNase pre-electrophoresis treatment.

**Figure 2:**
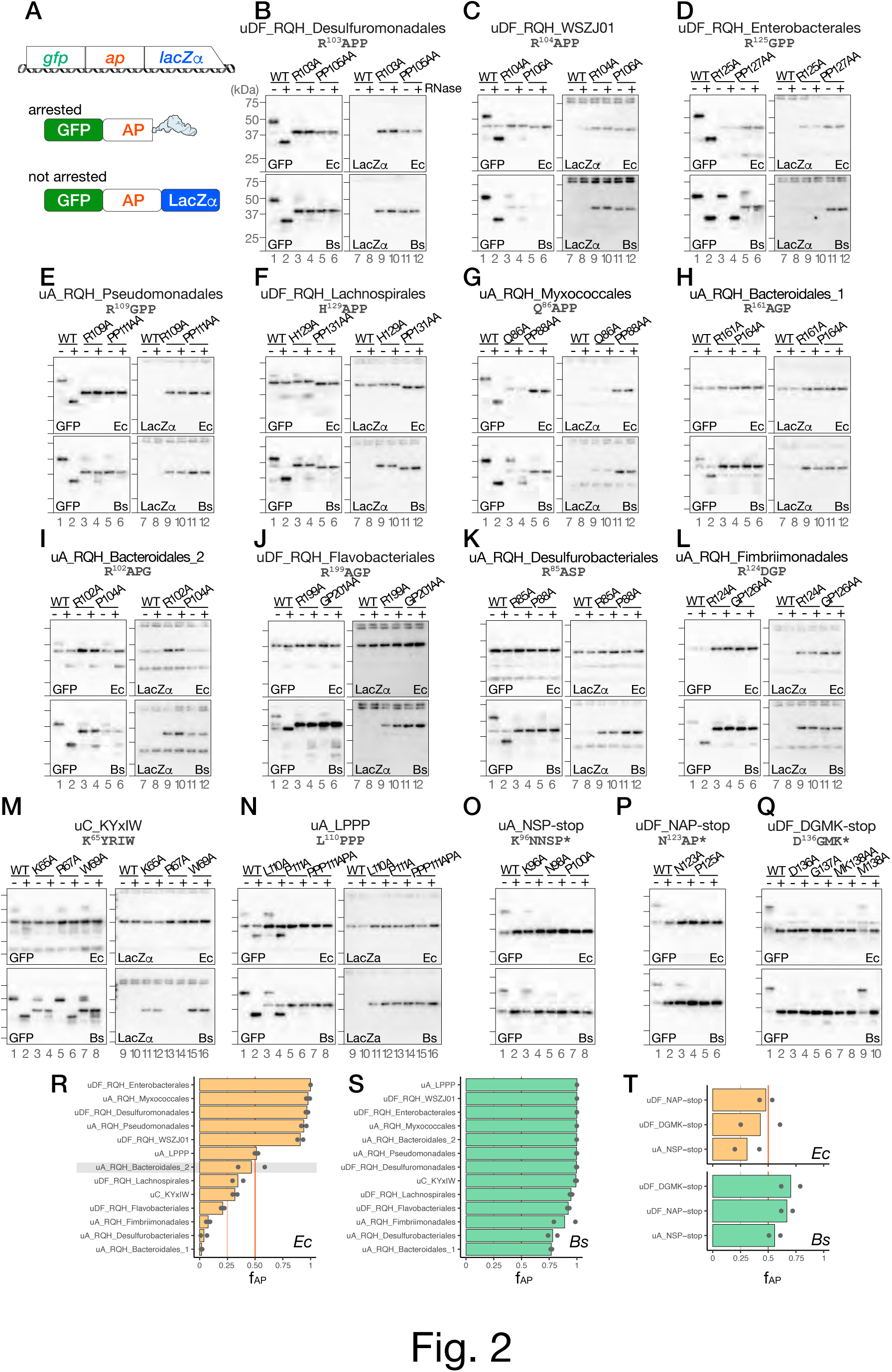
*In vitro* analysis of translation arrest. **a**, Schematic representation of the *gfp-ap-lacZα* translational fusion template used for *in vitro* coupled transcription/translation and its translation products. **b–q,** Western blot analysis of the *in vitro* translation products. The reporter genes harboring wild-type (WT) or mutant derivatives of the putative arrest peptides indicated at the top of the figure were translated in the *Ec* and *Bs* PURE systems. The products were separated in neutral-pH gels and immunoblotted using anti-GFP (left) or anti-LacZα (right) antibodies. Before the separation, a portion of the samples were treated with RNase A (lanes indicated as +), to degrade the tRNA moiety. Molecular size standards are indicated as horizontal lines on the left of each membrane; from top to bottom: 75, 50, 37, and 25 kDa, respectively. **r–t,** Fraction arrest peptide (f_AP_) of each reporter calculated according to the following formula: (arrest product) / (total translation products). The means of two independent experiments are indicated by the orange and green bars, which correspond to those obtained from experiments using the *Ec* (orange) or *Bs* (green) PURE systems, respectively. The dots indicate individual data points.

Strikingly, all of the candidate arrest peptides tested here stalled the ribosome in the *Bs* PURE system; moreover, many candidate arrest peptides also arrested translation in the *Ec* PURE system (Fig. 2). For instance, translation of the reporter for uDF_RQH_Desulfuromonadales in *Ec* PURE resulted in the accumulation of a major translation product of ∼50 kDa that was reactive to an anti-GFP, but not to an anti-LacZα, antibody (Fig. 2b, upper panels, lanes 1 and 7). The RNase pre-treatment resulted in a mobility shift (Fig. 2b, upper panel, lane 2), indicating that this was an arrested product. A minor full-length product of ∼43.5 kDa was also detected (lanes 1, and 2). The fraction arrest peptide (f_AP_), which indicates the proportion of the arrest product among the total translation products was 0.96, suggesting that almost all of the translating ribosomes were stalled (Fig. 2r). The replacement of conserved arginine (Arg_103_) or proline (Pro_105_-Pro) residues by alanine residue(s) led to the predominant accumulation of the full-length product (Fig. 2b, R103A and PP105AA, upper membranes). Similar results were obtained for translation using the *Bs* PURE system, which yielded an f_AP_ value of 1 (Fig. 2b, lower membranes, and 2s). Other RQH family members, i.e., uDF_RQH_WSZJ01, uDF_RQH_Enterobacterales, uA_RQH_Pseudomonadales, and uA_RQH_Myxococcales, which carry R_104_APP, R_125_GPP, R_109_GPP, and Q_86_APP sequences, respectively (Fig. 2c–e, g, r and s) arrested translation in both *Ec* and *Bs* PURE, with f_AP_ > 0.9.

The remaining six members of the RQH family (uDF_RQH_Lachnospirales, uDF_RQH_Flavobacteriales, uA_RQH_Fimbriimonadales, uA_RQH_Desulfurobacteriales, uA_RQH_Bacteroidales_1, and uA_RQH_Bacteroidales_2) arrested translation to lesser extents when translated using the *Ec* PURE system, whereas they efficiently arrested translation in the *Bs* PURE system (Fig. 2). Although uA_RQH_Bacteroidales_2 exhibited an f_AP_ value of 0.47 in the *Ec* PURE system, our toeprinting failed to identify the ribosome-stalling signal at the RAPG sequence (see below), which raised the possibility that the translation arrested product detected in the *Ec* PURE system was produced by ribosome stalling at a site other than the RAPG sequence. In accordance with this notion, neither the R102A nor the P104A mutation significantly abolished the translation arrest in *Ec* PURE, whereas they did so in *Bs* PURE (Fig. 2i).

In most cases, alanine substitution of the conserved arginine or proline(s) in the RQH family members abolished or compromised translation arrest (Fig. 2b–l), thus highlighting the general importance of these conserved residues. Similarly, the Q86A substitution in the QAPP motif of uA_RQH_Myxococcales significantly reduced the efficiency of the translation arrest in both the *Ec* and *Bs* PURE systems. In turn, the H129A substitution in the HAPP sequence of uDF_RQH_Lachnospirales compromised translation arrest in *Bs* PURE, whereas the minor translation arrest observed in the *Ec* PURE system was unaffected by this same mutation. In contrast, the R125A mutation in uDF_RQH_Enterobacterales did not abolish the translation arrest in the *Bs* PURE system, whereas it did so in the *Ec* PURE system (Fig. 2d, R125A).

### Novel sequences that cause translation arrest in vitro

Next, we focused on uORFs carrying C-terminal conserved motifs unrelated to known arrest peptides. Translation of the uC_KYxIW reporter using *Bs* but not *Ec* PURE resulted in the predominant accumulation of an RNase-sensitive arrest product (Fig. 2m, wild-type (WT)). The f_AP_ values obtained using *Ec* and *Bs* PURE were 0.32 and 0.99, respectively (Fig. 2r and s). The arrest was impaired by the substitution of the highly conserved Lys_65_ or Trp_69_ residue with alanine (Fig. 2m, K65A, and W69A, lower panels), but not by that of the less-conserved Arg_67_ residue (Fig. 2m, lower panels, R67A).

A homolog of uA_LPPP also efficiently stalled *B. subtilis* ribosomes *in vitro*, with an f_AP_ value of 1 (Fig. 2n, lower panels, and 2s). The fraction of the arrested product was reduced when the Leu_110_, Pro_111_ or Pro_111_/Pro_113_ was substituted with alanine(s) (Fig. 2n, lower panels). In contrast, uA_LPPP only modestly stalled *E. coli* ribosomes, with an f_AP_ value of 0.51 (Fig. 2n, upper panels, and 2r). The mutations of proline residues but not Lue_110_ decreased the accumulation of peptidyl-tRNA in *Ec* PURE (Fig. 2n, upper panels).

Regarding the candidate arrest peptides uA_NSP-stop, uDF_NAP-stop, and uDF_DGMK-stop, we constructed a series of *gfp-ap* reporters in which a gene segment encoding the C-terminal soluble region and the subsequent stop codon of the candidate arrest peptide were fused after the *gfp* gene. The f_AP_ was calculated based on the fraction of the peptidyl-tRNA among the total translation products. Approximately 30%–50% of the translation products of uA_NSP-stop, uDF_NAP-stop, and uDF_DGMK-stop were accumulated as a peptidyl-tRNA form when translated using *Ec* PURE (Fig. 2o–q, upper panels, WT, and 2t, upper graphs), whereas more than 50% of the translation products appeared as a peptidyl-tRNA form in *Bs* PURE (Fig. 2o–q, lower panels, WT, and 2t, lower graphs). In contrast, the direct fusion of the gene segment encoding the C-terminal region to *lacZa* led to the predominant accumulation of the full-length products (Supplementary Fig. 13), suggesting the importance of the termination codon for the arrest.

The alanine substitutions of Lys_96_, Asn_98_, and Pro_100_ in uA_NSP-stop reduced the peptidyl-tRNA accumulation in both *Ec* and *Bs* PURE (Fig. 2o), suggesting that the arrest depends on the amino acid residues or codons located at the −5,−3 and –1 positions from the C-terminal end (Fig. 2o, K96A, N98A, and P100A). Essentially similar results were obtained for uDF_NAP-stop and uDF_DGMK-stop; the stalling was affected by mutations of the residues located at positions −3 and −1 in the case of uDF_NAP-stop, and −4, −3, −2, and −1 in the case of uDF_DGMK-stop (Fig. 2p and q).

### Determination of ribosomal-stalling sites using a toeprinting assay

To determine the ribosome stalling site, we performed a toeprinting assay, in which we employed a fragment analysis to determine the size of the toeprint product (Fig. 3a)^33–35^. A control experiment confirmed that the toeprint product generated by the SecM-stalled ribosome appeared as a single peak, with its size indicating that the ribosome stalled with the P-site at the Gly_165_ (Fig. 3b and Supplementary Fig. 14), as demonstrated previously^34^. We then identified the stalling sites of the arrest peptides identified in this study using *Bs* PURE (Fig. 3c–f and Supplementary Fig. 15-30). For the arrest peptides that efficiently stalled both the *E. coli* and *B. subtilis* ribosomes, we also used the *E. coli* ribosome and confirmed that the estimated stalling site was identical. The results of our toeprinting analysis of the RQH family members revealed that, in all cases, the ribosome stalled when the codon for the third residue of the (R/Q/H)-(A/G/D)-(P/G/S)-(P/G) motif was in the P-site (Fig. 3c and d). Previous studies demonstrated that the stalling of ApcA, ApdA, and ApdP also occured at the third position of the RAPG/RAPP sequences, suggesting that these arrest peptides share a common underlying mechanism.

**Figure 3:**
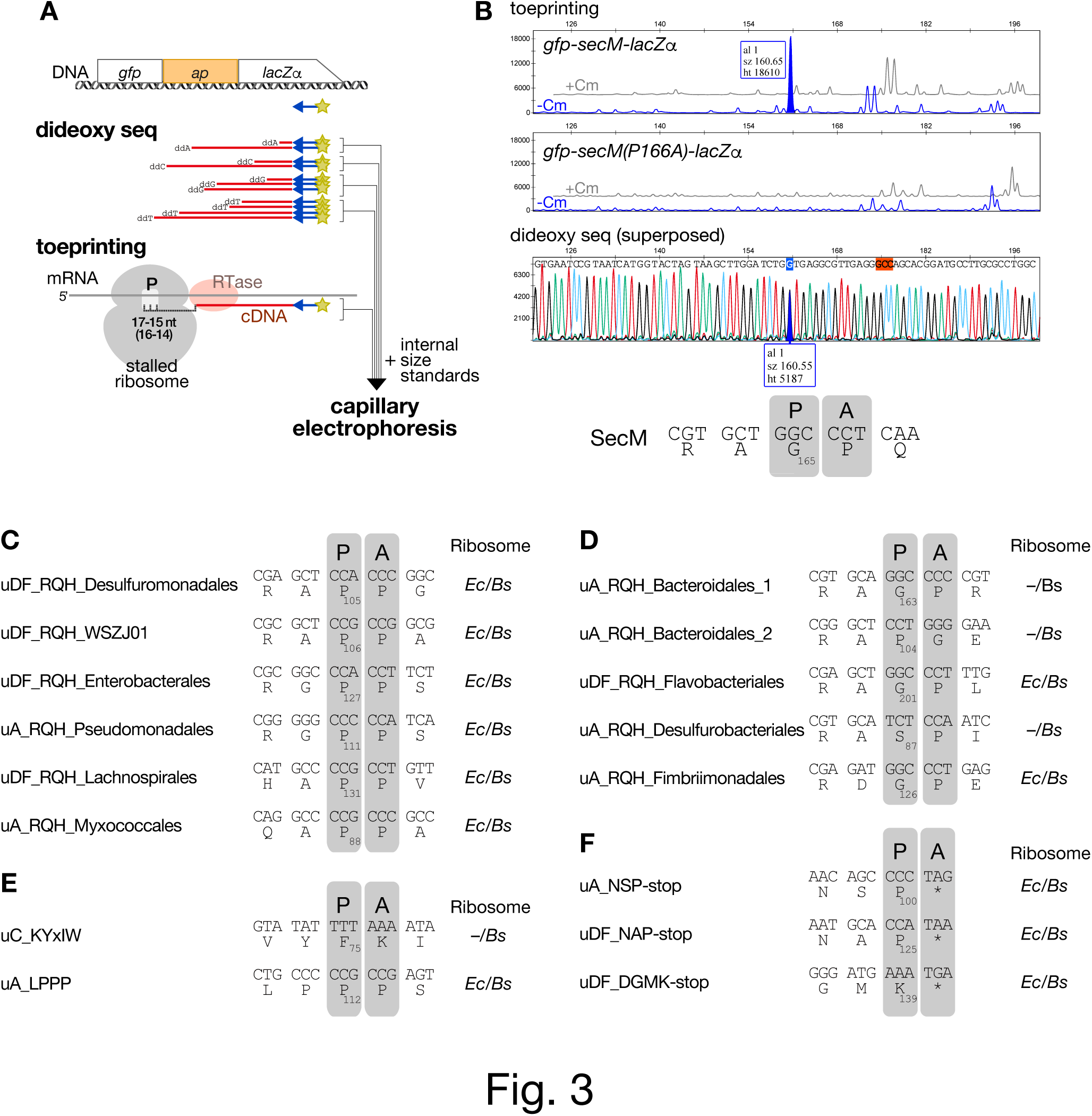
Determination of the arrest sites using a toeprinting analysis. **a**, Procedure used for toeprinting. **b,** Toeprinting of SecM. The gray (+Cm) and blue (+Cm) lines in the top and middle panels indicate the signals obtained from experiments performed in the presence or absence of chloramphenicol, respectively. The stalling-dependent toeprint signal specifically obtained from the experiment using the wild-type, but not in that using the arrest-defective P166A mutant, is filled in blue. The peak corresponding to the stalling-specific toeprint signal and its nucleotide in the dideoxy sequencing data (bottom panel) are filled and marked in blue, respectively. The estimated codon in the P-site of the stalled ribosome is marked in red. The estimated stalling site is indicated as the codons in the P-(P) and A-(A) sites of the stalled ribosome and the P-site codon number. Additional details and raw plots are provided in Supplemental Figure 14. **c–f,** Estimated ribosome-stalling sites of the candidate monitoring substrates. The stalling sites determined for either or both *E. coli* and *B. subtilis* ribosomes are shown. “P” and “A” represent P-site and A-site positions, respectively. Raw plots are provided in Supplemental Figures 15–30.

Interestingly, our toeprinting analysis revealed that the ribosome stalling of uC_KYxIW occurred at a site located at a more C-terminal position than the conserved motif. The toeprinting of uC_KYxIW produced four consecutive toeprint signals (Supplementary Fig. 26). The strongest signal indicated that the ribosomes stalled at the Phe_75_ codon in the P-site (Fig. 3e). Interestingly, the PTC-proximal residues were not well conserved among the homologs (Supplementary Fig. 3). The crucial Lys_65_ and Trp_69_ should be separated from the PTC by 11 and 7 residues, respectively, implying that they were situated in the mid-tunnel region in the NPET of the stalled ribosome (Fig. 2m). Further experiments will be necessary to address whether the minor signals indicate an additional stalling.

The toeprinting of uA_LPPP yielded two toeprint signals that indicated that the ribosome stalled when the P-site was at the third codon (Pro_112_) of the L_110_PPP motif (Fig. 3e). Toeprinting of both uA_NSP-stop and uDF_NAP-stop yielded signals indicating that the stalling occurred when the A-site was at the stop codon (Fig. 3f, Supplementary Fig. 28 and 29). In the case of uDF_DGMK-stop, the size of the major toeprint signal suggested that the stalling occurred either with the codon for Lys_139_ or the stop codon located at the A-site (Supplementary Fig. 30). The importance of the stop codon for the arrest of uDF_DGMK-stop (Supplementary Fig. 13) renders it more likely that uDF_DGMK-stop induces the arrest when the stop codon is at the A-site (Fig. 3f).

### In vivo reporter assay to determine the efficiency of the translation arrest

Next, we examined the efficiency of the elongation arrest *in vivo* using the *gfp-ap-lacZ* in-frame fusion reporters (Fig. 4a). Elongation arrest before *lacZ* will result in a low β-galactosidase activity. We compared the β-galactosidase activity of WT arrest peptides with those of arrest-defective mutant variants, and calculated the translation-arrest index (TAI), which is the ratio of the β-galactosidase activity of a mutant to that of WT (Fig. 4b–i and Supplementary Fig. 31). A high TAI value indicates an efficient translation arrest *in vivo*, whereas a TAI value as low as 1 suggests the absence of appreciable translation arrest *in vivo*. For example, *E. coli* cells harboring the reporter of uDF_RQH_Desulfuromonadales exhibited a low level of β-galactosidase activity (187 U), whereas the activities of the R103A and PP105AA mutant derivatives were 2,233 and 2,333 U, respectively (Fig. 4b). The TAI value calculated based on the β-galactosidase activities of the WT and PP105AA mutant strains was 12.51, which was indicative of an efficient translation arrest in *E. coli* (Fig. 4i and Supplementary Fig. 31). A similar result was obtained in the case of expression in *B. subtilis*, albeit with a relatively low TAI value of 3.18 (Fig. 4i and Supplementary Fig. 31).

**Figure 4:**
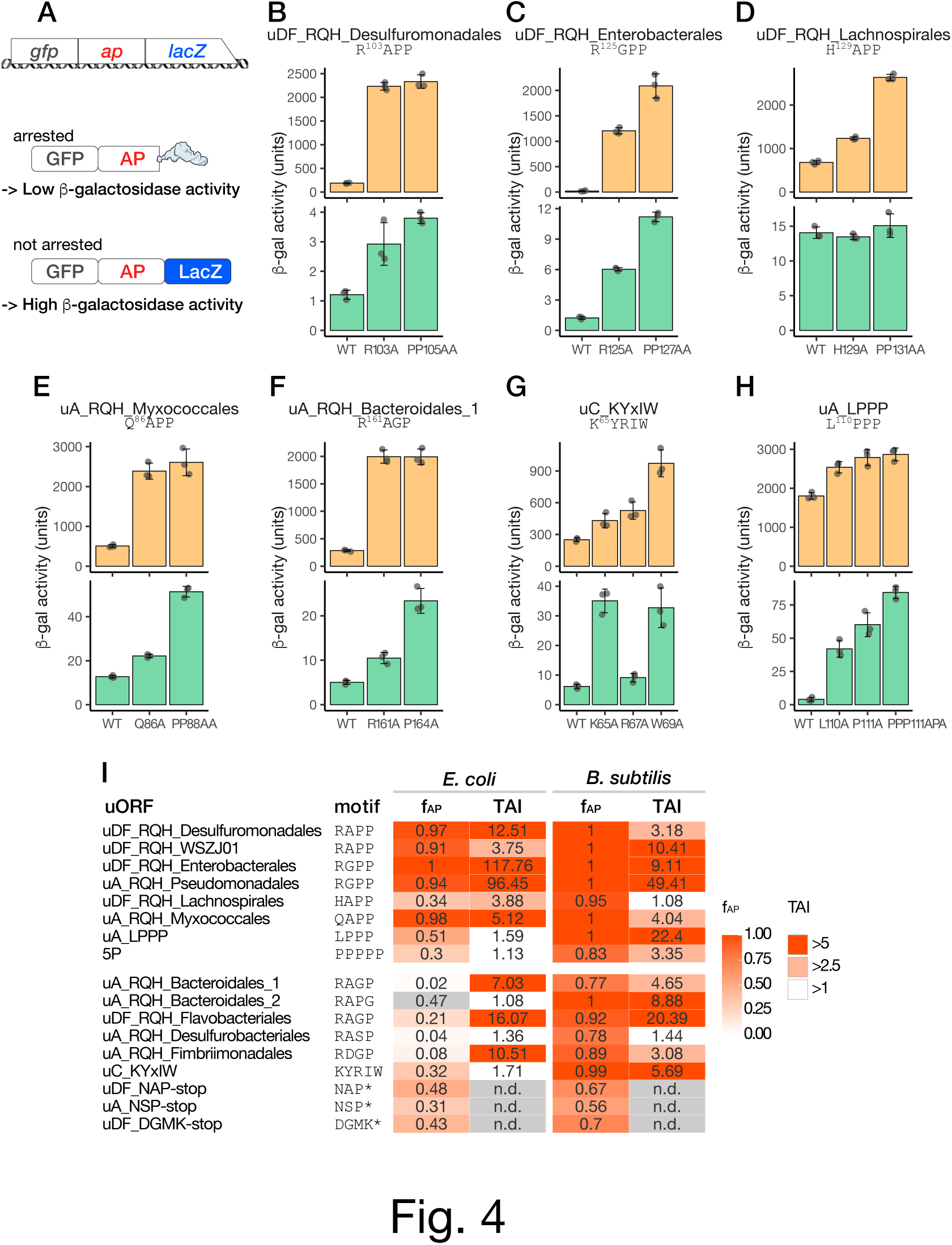
*In vivo* translation-arrest assay. **a**, Schematic representation of the reporter used for the *in vivo* assay and its translation products. **b–h,** β-galactosidase activity (mean, n = 3) of *E. coli* (orange bars) and *B. subtilis* (green bars) cells harboring wild-type (WT) or mutant derivatives of the arrest peptide reporters. The error bars and dots represent standard deviations and individual data points, respectively. **i,** Summary of the *in vitro* and *in vivo* analyses of translation arrest. The translation arrest indexes (TAI) were calculated based on the *in vivo* β-galactosidase activities (mutant / wild-type) and are listed together with f_AP_ (related to Figure 2).

When expressed in *E. coli*, most of the RQH family members yielded a high TAI value, with the exception of uA_RQH_Desulfurobacteriales and uA_RQH_Bacteroidales_2 (Fig. 4i and Supplementary Fig. 31). The former was consistent with its low arrest efficiency in *Ec* PURE. As mentioned above, the stalling of the latter in *Ec* PURE may have occurred at a site other than the RAPP-like sequence. Such an unrelated stalling may occur only *in vitro*. Low TAI values were obtained for uDF_RQH_Lachnospirales and uA_RQH_Desulfurobacteriales from experiments using *B. subtilis* (Fig. 4i and Supplementary Fig. 31). We cannot provide a clear explanation for the apparent inconsistency between the results obtained *in vivo* and *in vitro*. Nevertheless, the high TAI values obtained for most of the RQH family members in *E. coli* and *B. subtilis* demonstrated that they arrest translation efficiently *in vivo*.

The TAI values of uA_LPPP in *E. coli* and *B. subtilis* were 1.59 and 22.4, respectively (Fig. 4i and Supplementary Fig. 31), suggesting that it efficiently stalls the *B. subtilis* but not *E. coli* ribosome *in vivo*. It is conceivable that the arrest of uA_LPPP observed in the *Ec* PURE system was an *in vitro* artifact caused by the absence of EF-P. The results of the *in vivo* analysis of uC_KYxIW showed a good agreement with those of its *in vitro* analysis, in which it stalled the *B. subtilis*, but not the *E. coli*, ribosome efficiently (Fig. 2).

### The translation arrest of uDF_RQH_Enterobacterales induces the downstream secDF gene

The translation arrest of SecM, MifM, and VemP results in the induction of downstream target genes^9–11,24^. A similar role could be expected for the newly identified arrest peptides. To test this possibility, we focused on a homolog of uDF_RQH_Enterobacterales derived from *Alteromonas naphthalenivorans*, which belongs to the same order (Enterobacterales) as *E. coli* (Supplementary Fig. 2). We identified two predicted stem-loop structures, i.e., stem-loop 1 and stem-loop 2 (Fig. 5a), the latter of which partially masked the SD sequence of the downstream *secDF2* gene. We hypothesized that the stalled ribosome will disrupt stem-loop 2, keep the SD sequence exposed and thereby allow the expression of the downstream *secDF2* gene. To test this hypothesis, we constructed a reporter in which the coding region of GFP-uDF_RQH_Enterobacterales (residues 43–131) was followed by the intergenic region and the *secDF2’-lacZ* reporter (Fig. 5a).

**Figure 5:**
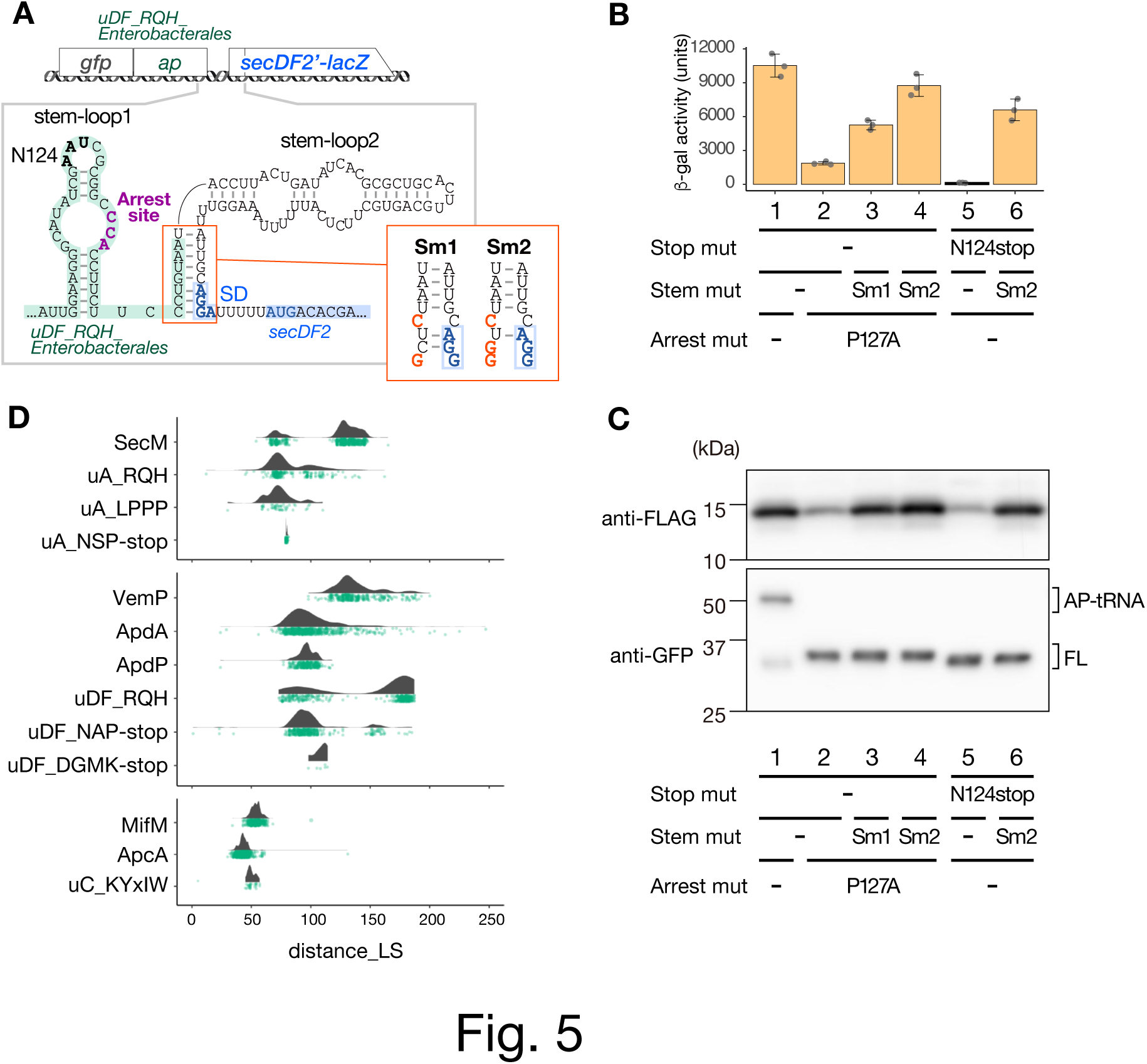
Arrest-dependent regulation of downstream genes. **a**, Schematic representation of the *lacZ* reporter used to examine the expression of a downstream gene (upper) and the secondary structures of the intergenic region located between *uDF_RQH_Enterobacterales* (green) and *secDF2* (blue) (bottom). The translation-arrest site is indicated in purple, and the N124 codon replaced by a stop codon within the mutant reporter in (B) and (C) is indicated in bold. The red panel shows the sequences of the Sm1 and Sm2 mutant reporters, in which the mutations introduced are indicated in bold red characters, whereas the putative SD sequence of *secDF2* is indicated in bold blue characters. **b,** *In vivo* analysis of *secDF2* regulation by uDF_RQH_Enterobacterales. Reporters harboring wild-type (lane 1) and a stop-codon-substitution mutant (lanes 5 and 6), as well as an arrest-defective P127A mutant (lane 2) and its derivatives with stem-mutations (lanes 3 and 4) of uDF_RQH_Enterobacterales, were expressed in *E. coli*, and the β-galactosidase activities were measured (means ± standard deviations, n = 3). **c,** *In vitro* reconstitution of the arrest-dependent induction of the downstream gene. LacZα-3xFLAG was used as the downstream reporter gene. The gene encoding SecDF2’-LacZa-3xFLAG carrying the upstream *gfp-uDF_RQH_Enterobacterales* was translated in the *Ec* PURE system, and the translation products were analyzed by Western blotting using anti-FLAG (upper panel) and anti-GFP (lower panel) antibodies, respectively. **d,** Raincloud plots of the distances between the arrest sites (P-site residues) and the last residues of the N-terminal localization signal (distance_LS).

An *E. coli* strain expressing the *secDF2’-lacZ* reporter described above exhibited a high β-galactosidase activity (10,531 units), which was drastically diminished by approximately 5.6-fold after the introduction of the P127A mutation in the R_125_GPP motif of uDF_RQH_Enterobacterales (Fig. 5b, columns 1, 2). Abolishment of the arrest by the P127A mutation was confirmed using *Ec* PURE (Fig. 5c, lower panel, lanes 1 and 2). These results suggest that the translation arrest of uDF_RQH_Enterobacterales strongly induces the expression of the downstream *secDF2* gene. The decreased induction of *secDF2* by the P127A mutation was partially counteracted by the disruption of two G-C base pairs (Sm1) or strongly counteracted by the disruption of three G-C base pairs (Sm2) in stem-loop 2 (Fig. 5a, b). These findings are consistent with the notion that the formation of the stem-loop 2 sequesters the SD sequence of the *secDF2* gene, thus repressing the induction of the *secDF2* gene. The premature translation termination that occurred after the introduction of a nonsense mutation at the 124^th^ codon (Fig. 5a) resulted in an even lower β-galactosidase activity compared with the basal activity observed for the P127A mutant; moreover, the abolishment of the induction was again counteracted by the Sm2 mutation (Fig. 5b, columns 5 and 6). We assume that the ribosome that translates the P127A derivative can still transiently disrupt the stem-loop 2, which is completely eliminated by the premature translation termination by the N124stop mutation.

The arrest-dependent induction of the downstream gene was also recapitulated *in vitro*, in which the translation arrest of the GFP-uDF_RQH_Enterobacterales derivatives (Fig. 5c, lower) and induction of its downstream gene (*secDF2-lacZα-3xFLAG* reporter; Fig. 5c, upper) were assessed by anti-GFP and anti-FLAG antibodies, respectively. Taken together, these data support the notion that the translation arrest of uDF_RQH_Enterobacterales is responsible for *secDF2* expression via a mechanism similar to that of SecM, MifM, and VemP.

### Bioinformatics analysis of the length of the spacer between the protein localization signal and the arrest site

For an arrest peptide to function as a Sec– or YidC-monitoring substrate, the translation arrest must be released in a secretion– or membrane-insertion-dependent manner^27^. An optimal distance between the N-terminal TM segment and the arrest site is crucial for the membrane-insertion-dependent arrest cancellation if the TM segment adopts the type I (N-out/C-in) orientation^36,37^, which is a topology that is often observed for YidC substrates^38^. Conversely, this distance can vary without impairing the localization-dependent arrest release if the N-terminal localization signal is either the Sec-dependent secretion signal or type II (N-in/C-out) TM segment^37^. We envisioned that this trend might be shared by known monitoring substrates, i.e., SecM, MifM, and VemP, as well as by other arrest peptides if they actually function as monitoring substrates.

To test these possibilities, we first determined the distance between the putative N-terminal localization signal and the arrest site of each homolog of SecM and plotted them to generate a distribution diagram (Fig. 5d, upper). We identified two discrete peaks with median distances of 131.5 (494 genomes) and 71 (145 genomes), respectively, suggesting that SecM homologs can be divided into two classes, i.e., long and short variants, as reported previously^39^. In contrast, VemP, which is another known Sec-monitoring substrate, exhibited relatively long spacer lengths between the N-terminal localization signal and the arrest site, with a median distance of 133 (Fig. 5d, middle). In contrast, MifM, which is a YidC-monitoring substrate, exhibited a relatively narrow distribution pattern, with a shorter median distance of 54. These observations were consistent with our assumption that the spacer length of the YidC-monitoring substrate is relatively shorter and less variable than that of the Sec-monitoring substrate.

We performed a similar analysis for ApcA, ApdA, ApdP, and arrest peptides identified in this study. Our analysis revealed that the arrest peptides encoded by the uORFs of *secA* (Fig. 5d, upper) or *secDF* (Fig. 5d, middle) typically had relatively longer spacer regions, with a wide range of length variations observed in most cases compared with MifM, ApcA, and uY_KyxIW, which are encoded by the uORF of *yidC* (Fig. 5d, lower). These observations suggest that these arrest peptides also share the molecular feature expected of monitoring substrates.

### Translation arrest of RAPP/RAGP motif-containing proteins in E. coli and B. subtilis

Previous and current studies revealed that many of the arrest peptides encoded by the uORFs of the *sec* or *yidC* gene harbor RAPP/RAGP-like motifs. We envisioned that other RAPP-containing proteins encoded by ORFs unrelated to the *sec* or *yidC* gene might also stall the ribosome. Therefore, we searched for proteins containing RAPP, RGPP, HAPP, HGPP, QAPP, QGPP, RAGP, RAPG, and RPPP sequences in the *E. coli* and *B. subtilis* proteomes (Supplementary Table 5). Among them, we chose 14 and 9 RAPP-like sequences derived from *E. coli* and *B. subtilis*, respectively, to examine their capability of stalling the ribosomes *in vitro*. The gene fragment encoding the RAPP-like motif was cloned between the *gfp* and *lacZ* genes (Fig. 6a). Translation using *Ec* or *Bs* PURE revealed that most of them did not arrest translation efficiently (Supplementary Fig. 32). However, we found that a sequence derived from *B. subtilis* YwcI, i.e., a short peptide of 100 amino acids with an RAGP motif at residues 88–91, efficiently induced translation arrest *in vitro* (Fig. 6b), with an f_AP_ value of 0.95. This arrest was abolished by replacing Pro_91_ with alanine (Fig. 6b, P91A), suggesting that the proline of the RAGP motif is crucial for the arrest. A toeprinting analysis demonstrated that the ribosome stalled with the P-site at the Gly_90_ codon (Fig. 6c and Supplementary Fig. 33). Finally, an *in vitro* reporter assay revealed a low level of β-galactosidase activity for the WT *ywcI* reporter, which was elevated after the introduction of the arrest-defective P91A mutation, resulting in a TAI value of 5.7. These results demonstrated that YwcI is a novel arrest peptide derived from *B. subtilis*.

**Figure 6:**
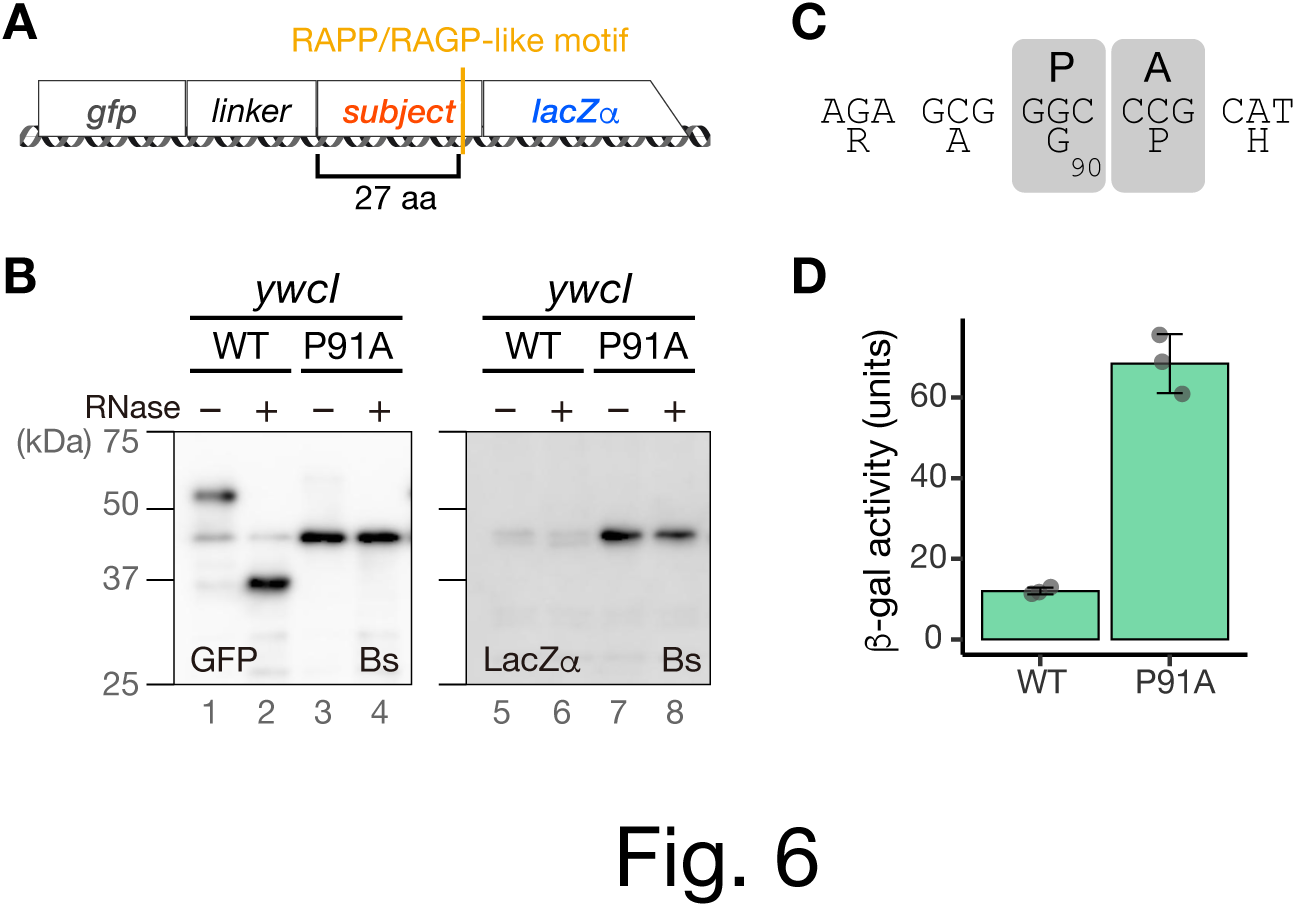
Translation arrest by *B. subtilis* YwcI. **a**, Schematic representation of the *lacZα* reporter used for the *in vitro* translation of *E. coli* and *B. subtilis* proteins containing the RAPP/RAGP-like motif. The gene segments for target proteins containing RAPP-like sequences were sandwich-fused between *gfp* with a linker and *lacZα*. **b,** *In vitro* analysis of the translation arrest of YwcI. Fusion genes harboring wild-type (WT) or the P91A mutant derivatives of *ywcI* were translated using the *Bs* PURE system. The products were separated on neutral-pH gels and immunoblotted using anti-GFP (left) or anti-LacZα (right) antibodies. Before the separation, a portion of the samples were treated with RNase A (lanes indicated as +), to degrade the tRNA moiety. **c,** Estimated ribosome stalling site of *ywcI*. “P” and “A” represent the P-site and A-site positions, respectively. Raw plots are provided in Supplemental Figure 34. **d,** *In vivo* translation-arrest analysis. β-galactosidase activity (means, n = 3) of *B. subtilis* cells harboring wild-type (WT) or P91A mutant reporters. The error bars and dots represent standard deviations and individual data points, respectively.

## Discussion

Our extensive screening led to the identification of dozens of arrest peptides that could possibly serve as Sec or YidC-monitoring substrates. Those include more than 10 members of the RQH family, which bear motifs similar to one another and to some of the arrest peptides identified previously, i.e., SecM, ApcA, ApdA, and ApdP. Other arrest peptides that bear novel arrest-inducing sequences were also identified. Our results highlight a unique pattern of evolution of the bacterial arrest peptides encoded upstream of the *sec* or *yidC* genes, which likely have emerged repeatedly in various bacterial species, resulting in a patchy and widespread phylogenetic distribution.

Arrest peptides often stall the ribosome in a species-specific manner, and, consistent with this notion, they generally have a narrow phylogenetic distribution. Therefore, the occurrence of the RQH family members in various bacterial phyla is an unusual evolutionary trait that has not been observed for other arrest peptides. Furthermore, we identified YwcI, an RAGP-containing arrest peptide encoded by a gene upstream of *sacT* (Fig. 6). SacT is an anti-transcriptional terminator of the *sacP* and *sacA* genes, which encode a sucrase and sucrose-specific permease, respectively^40–42^. This result suggests that the function of RAPP-containing arrest peptides is not limited to the regulation of the Sec or YidC pathway. In accordance with this notion, recent studies have suggested that the CruR and CutF uORFs, which harbor RAPP and polyproline sequences, respectively, play roles in regulating the downstream ORFs encoding the TonB-dependent transporter BfrG and multicopper oxidase CutO, respectively^43,44^, although the translation arrest of either of them has yet to be demonstrated experimentally.

The sharing of a similar RAPP-like sequence makes it plausible that these arrest peptides employ a similar nascent-chain–ribosome interaction near the PTC to achieve ribosome stalling. This might also be the case for the arrest-inducing sequences containing RAPP-like motifs that were identified in a screening using a random sequence library^45,46^. However, the stalling efficiencies on *E. coli* and *B. subtilis* ribosomes varied among each arrest peptide (Fig. 2 and 4), suggesting that the residues located upstream of the RAPP motif play an important role in determining the species specificity, as suggested previously for ApdA, and ApdP^28^. In addition, during the identification of YwcI, we found that most of the *E. coli* and *B. subtilis* proteins that contained RAPP-like motifs did not stall the ribosome (Supplementary Fig. 32). These observations point to the importance of the N-terminal region in the stalling ability. The N-terminal region may interact with the mid-tunnel region, in which ribosomal proteins, uL4 and uL22, as well as ribosomal RNA components sometimes encompass species-specific structures^47,48^. This might have resulted in diverse and, sometimes, species-specific N-terminal sequences.

A key question arising from the current results is why the RAPP-containing arrest peptide alone was prevalent across the bacterial domain. One possibility is that the RAPP-like sequence serves as a versatile “seed” to tailor-make arrest-inducing sequences that can be flexibly optimized beyond the species-specific structural differences of each ribosome. The lack of an appreciable sequence similarity among the N-terminal region of the RAPP-containing arrest peptides suggests that a wide variety of N-terminal sequences might be compatible with the RAPP-like sequence without disrupting the ribosome-stalling capability, as suggested for SecM^49^.

Another intriguing question is whether all of the arrest peptides that contained the RAPP-like sequence were derived from a common evolutionary origin or emerged repeatedly in different species and evolved independently. Although it is difficult to rule out one of these two possibilities, we favor the latter scenario for the following reasons: (i) these arrest peptides lacked overall sequence similarity to one another, with the exception of the RAPP-like motif; (ii) gene contexts were divergent among these arrest peptides; and (iii) the repetitive occurrence of such a short sequence motif during evolution seems possible.

We also successfully identified arrest peptides that bore novel arrest motifs. Those included uC_KYxIW, uA_LPPP, uA_NSP-stop, uDF_NAP-stop, and uDF_DGMK-stop. Among them, uC_KYxIW was unique in that its conserved and arrest-essential motif was located 7–11 residues distal from the stalling site and the PTC-proximal residues were less conserved (Supplementary Fig. 3). The identification of various distinct arrest sequences suggests that bacteria have evolved various arrest-inducing mechanisms and that there must be more unidentified arrest-inducing sequences.

In the present study, the results of the *in vivo* and *in vitro* translation arrest assays were generally in great agreement with each other. However, inconsistencies remained in some cases. For instance, uDF_RQH_Lachnospirales, uA_RQH_Myxococcales, uA_RQH_Desulfurobacteriales, and uA_RQH_Fimbriimonadales stalled the *B. subtilis* ribosomes efficiently *in vitro*, but exhibited a relatively lower arrest efficiency in the *in vivo* experiment (Fig. 4i). One possible explanation for this result is that the ribosome stalling caused by the foreign peptides was somehow subjected to release by a cellular factor, resulting in a reduced stability of arrest *in vivo*. Another possibility is that the translation product of the reporter became a target of proteolysis *in vivo*, thus causing an apparent inconsistency. Conversely, uA_RQH_Fimbriimonadales efficiently stalled the *E. coli* ribosome *in vivo* but not *in vitro*. It is formally possible that an *in vivo* co-factor or specific condition is required for the stabilization of the arrest.

We demonstrated that the elongation arrest by *A. naphthalenivorans* uDF_RQH_Enterobacterales led to the induction of the downstream *secDF2* gene (Fig. 5a–c). The bioinformatics analysis of the spacer length between the localization signal and the arrest motif also suggest that the arrest peptides identified in this study share the molecular feature expected of monitoring substrates (Fig. 5d). These observations support the notion that the arrest peptides identified in this study function as monitoring substrates.

This study opened up the possibility that the search for proteins containing the RAPP-like sequence may allow the identification of novel arrest peptides, as demonstrated for *B. subtilis* YwcI. Further identification and characterization of novel arrest peptides will provide insights into the shared or lineage-specific evolution of arrest peptides, during which each bacterium must have achieved various physiological functions and mechanisms of translation regulation through common or unique interactions between the ribosome and the nascent peptide chains. The exploration of various regulatory arrest peptides will unveil unidentified principles via which the ribosome translates genetic information into cellular functions in a manner that is beyond our current understanding.

## Materials and Methods

### Bioinformatics screening of putative monitoring substrates

The *in silico* screening for ORFs encoding monitoring-substrate-like arrest peptides was conducted as described below and detailed in the Supplementary Procedures. A total of 30,175 representative bacterial genome sequences were obtained from the GTDB^29^ release 202 (Supplementary Table S1). The *secA*, *secDF*, and *yidC* genes were identified from each bacterial genome using BLAST+^50,51^ version 2.13.0. Subsequently, their uORFs encoding small secretory or transmembrane proteins (<300 aa) of unknown function (annotated as hypothetical, putative, uncharacterized, unknown, DUF, membrane protein, or extracytoplasmic protein) or homologs of known monitoring substrates (SecM, MifM, and VemP) were identified as described in the Supplementary Procedures. Putative secretion signals and TM segments were predicted using SignalP^52^ version 6.0, deepTMHMM^53^ version 1.0.18, and TMHMM^54^ version 2.0. These uORFs were then subjected to a clustering analysis using MMseqs2^30^ version 14-7e284, followed by alignment with each cluster using MAFFT^55,56^ version 7.490, to identify homology groups with C-terminal sequences shared within each homology group. This analysis was conducted using R scripts developed in-house (with R version 4.0 or later).

### Phylogenetic tree

Phylogenetic tree data were downloaded from the GTDB (release 202). If needed, the tree was split into a specific phylum using the “drop.tip” function of ape^57^ version 5.6.2. The tree was visualized and decorated using iTol^58^ version 6.

### Bacterial strains and plasmids

The *B. subtilis* strains, plasmids, and DNA oligonucleotides used in this study are listed in Tables S6, S7, and S8, respectively. The preparation of synthetic DNAs for candidate monitoring substrates was outsourced (Thermo Fisher). Plasmids were constructed via standard cloning methods, including PCR using PrimeSTAR GXL (Takara), and DpnI treatment (Takara); moreover, Gibson assembly^59^. Sera-Mag Carboxylate-Modified Magnetic Particles (Cytiva, 65152105050250) was used to purify double-stranded DNA^60^ and sequencing products^61^. The *B. subtilis* strains were constructed by transformation involving double homologous recombination between chromosomal DNA and the plasmids introduced into *B. subtilis* competent cells. The resulting recombinant clones were validated based on their antibiotic-resistance markers.

### Culture media and growth conditions

*B. subtilis* cells were cultured in LB medium. *E. coli* cells were cultured in LB medium supplemented with 100 μg/ml ampicillin. Cells were cultured at 37°C and collected for Western blotting or β-galactosidase activity assay when they reached an optical density of 0.5–1.0 at 600 nm (OD_600_).

### Antiserum production

The production of the anti-LacZα antiserum was outsourced to Eurofins. Two chemically synthesized peptides, NH_2_-CRNSEEARTDRPSQQ-COOH and NH_2_-CTDRPSQQLRSLNGE-COOH, corresponding to residues 38–51 and 45–58 of *E. coli* LacZ, respectively, were used for immunization. N-terminal cysteines were added to both polypeptides, for their conjugation to the carrier protein Keyhole limpet hemocyanin.

### In vitro translation and Western blotting

Bacterial reconstituted transcription–translation coupling systems^31,32^ were used in the *in vitro* translation assay. Specifically, for *in vitro* translation using *E. coli* ribosomes, we utilized PUREfrex version 1.0 (GeneFrontier) according to the manufacturer’s protocol. For *Bs* PURE, purified *B. subtilis* ribosomes were used at the final concentration of 1 μM in the PURE system, without adding *E. coli* ribosomes. Then, 2.5 U/μL of T7 RNA polymerase (Takara) was added, to ensure transcription. The *in vitro* translation reaction was primed using the DNA templates listed in Table S8. The translation reaction was carried out for 30 min at 37°C and was stopped by adding 2× SDS–PAGE loading buffer, for Western blotting. A portion of the sample was further treated with 0.2 mg/ml RNase A (Promega) at 37°C for 10 min, to degrade the tRNA moiety of the peptidyl-tRNA, if necessary. The translation products were separated on a 10% polyacrylamide gel that was prepared using WIDE RANGE Gel buffer (Nacalai Tesque), according to the manufacturer’s instructions, then transferred onto a PVDF membrane (Merck, IPVH00010) and subjected to immune-detection using antibodies against GFP (mFX75; Wako) or LacZα. Images were acquired and analyzed using an Amersham Imager 600 luminoimager (GE Healthcare), and the band intensity was quantified using ImageQuant TL (GE Healthcare).

### Toeprinting assay

*In vitro* translation was carried out using the *Ec* PURE or *Bs* PURE system at 37°C for 20 min in the presence or absence of 0.1 mg/mL chloramphenicol, a translation inhibitor. The translation reaction mixture was then mixed with the same volume of the reverse transcription mixture containing 50 mM HEPES-KOH, pH7.6, 100 mM potassium glutamate, 2 mM spermidine, 13 mM magnesium acetate, 1 mM DTT, 2 μM of oligonucleotide labeled with 6-carboxyfluorescein (6-FAM) at the 5’ end (5’– AACGACGGCCAGTGAATCCGTAATCATGGT–3’, Invitrogen), 50 μM each dNTP, and 10 U/μL ReverTra Ace (Toyobo), then incubated further at 37°C for 15 min. The reaction mixture was diluted 5-fold with the NTC buffer (Macherey-Nagel), and the reverse transcription products were purified using a NucleoSpin Gel and PCR Clean-up kit (Macherey-Nagel). The reverse transcription products were eluted with 30 μL of HiDi formamide (Thermo Fisher). Samples were then mixed with 10 μL of 10-fold-diluted GeneScan 500 LIZ dye size standard (Thermo Fisher, 4322682), then heated at 96°C for 3 min just before capillary electrophoresis. The dideoxy DNA samples used as size markers for sequencing were prepared using a Thermo Sequenase Dye Primer Manual Cycle Sequencing Kit (Thermo, 79260), Thermo Sequenase Cycle Sequencing Kit (Thermo, 785001KT), or Thermo Sequenase DNA Polymerase (Cytiva, E79000Y), according to the manufacturer’s instruction, with some modifications. The DNA polymerase reaction was carried out using the same sets of template DNA and primer used for the toeprint assay. Each reaction mixture contained 0.44 μM of the 6-FAM-labeled primer, 60 μM each deoxynucleotide triphosphate (dATP, dCTP, dGTP, and dUTP), and 0.6 μM dideoxynucleotide triphosphate (either ddATP, ddCTP, ddGTP, or ddUTP). The sequencing products were purified using Sera-Mag speed beads and eluted with HiDi formamide. Next, 2 μL of a 10-fold-diluted GeneScan 500 LIZ dye size standard was added. If needed, the toeprinting product was further diluted before electrophoresis using HiDi formamide. The toeprinting and dideoxy sequencing products were then subjected to fragment analysis on a Seqstudio genetic analyzer (Thermo Fisher). Fragment data were analyzed and visualized using the GeneMapper software version 6 (Applied Biosystems), and processed further using Adobe Illustrator. The signals obtained from dideoxy sequencing were colored green (A), blue (C), black (G), and red (T), and then superposed for presentation (Fig. 3b and Supplementary Fig. 14).

### In vivo β-galactosidase assay

The β-galactosidase assay was performed as described previously^6^. A 100-μL aliquot of the culture was transferred to a well in a 96-well plate, and OD_600_ was recorded. We mixed the culture with 50 μL of Y-PER reagent (Thermo Fisher) for 20 min at room temperature, to disrupt the cells. In the case of *E. coli* cells, the mixture was diluted 10-fold and further subjected to freeze–thaw treatment, to ensure cell disruption. Subsequently, 30 μL of *O*-nitrophenyl-β-D-galactopyranoside (ONPG) in Z-buffer (60 mM Na_2_HPO_4_, 40 mM NaH_2_PO_4_, 10 mM KCl, 1 mM MgSO_4_, and 38 mM β-mercaptoethanol) was added to the cell lysate, and the OD_420_ and OD_550_ were measured at 28°C every 5 min over a period of 60 min. Arbitrary units of β-galactosidase activity were calculated using the following formula: [(1000 × V_420_ − 1.3 × V_550_) / OD_600_] for *B. subtilis*, or [10 × (1000 × V_420_ + 1.3 × V_550_) / OD_600_] for *E. coli*; where V_420_ and V_550_ are the first-order rate constants, OD_420_/min and OD_550_/min, respectively.

## Author Contributions

K.F., T.J., M.Y., and S.C. designed the research, K.F., T.J., M.Y., and S.C. performed experiments, K.F. performed bioinformatic analyses, K.F., H.T., and S.C. supervised the work, all authors analyzed the data, and K.F. and S.C. wrote the manuscript.

## Declarations of interest

none.

## Acknowledgments

We thank Machiko Murata and Naoko Muraki for their technical support. This work was supported by JSPS Grant-in-Aid for Scientific Research (Grant No. 16H04788, 26116008, 20H05926, and 21K06053 to SC, 19K16044, and 21K15020 to KF, 23K05017 for HT) and by JST, ACT X (Grant No. JP1159335 to H.T.)

## Declaration of Generative AI and AI-assisted technologies in the writing process

While preparing this work, the authors used DeepL and ChatGPT for English language editing. After using this tool, the authors reviewed and edited the content as needed and take full responsibility for the content of the publication.

## Notes

### Competing Interest Statement

The authors have declared no competing interest.

### Summary of Updates

Data not mentioned in the text were deleted from Figure 4.

## References

1. Ito, K. & Chiba, S. Arrest peptides: cis-acting modulators of translation. Annu. Rev. Biochem. 82, 171–202 (2013).

2. Dever, T. E., Ivanov, I. P. & Sachs, M. S. Conserved Upstream Open Reading Frame Nascent Peptides That Control Translation. Annu. Rev. Genet. 54, 237–264 (2020).

3. Chiba, S., Fujiwara, K., Chadani, Y. & Taguchi, H. Nascent chain-mediated translation regulation in bacteria: translation arrest and intrinsic ribosome destabilization. J. Biochem. 173, 227–236 (2023).

4. Wilson, D. N., Arenz, S. & Beckmann, R. Translation regulation via nascent polypeptide-mediated ribosome stalling. Curr. Opin. Struct. Biol. 37, 123–33 (2016).

5. Yang, Z., Iizuka, R. & Funatsu, T. Nascent SecM chain outside the ribosome reinforces translation arrest. PLoS One 10, e0122017 (2015).

6. Fujiwara, K., Ito, K. & Chiba, S. MifM-instructed translation arrest involves nascent chain interactions with the exterior as well as the interior of the ribosome. Sci. Rep. 8, 10311 (2018).

7. Su, T. et al. The force-sensing peptide VemP employs extreme compaction and secondary structure formation to induce ribosomal stalling. Elife 6, e25642 (2017).

8. Shanmuganathan, V. et al. Structural and mutational analysis of the ribosome-arresting human XBP1u. Elife 8, e46267 (2019).

9. Murakami, A., Nakatogawa, H. & Ito, K. Translation arrest of SecM is essential for the basal and regulated expression of SecA. Proc. Natl. Acad. Sci. U. S. A. 101, 12330–5 (2004).

10. Chiba, S., Lamsa, A. & Pogliano, K. A ribosome-nascent chain sensor of membrane protein biogenesis in Bacillus subtilis. EMBO J. 28, 3461–3475 (2009).

11. Ishii, E., et al. Nascent chain-monitored remodeling of the Sec machinery for salinity adaptation of marine bacteria. Proc. Natl. Acad. Sci. 112, E5513–E5522 (2015).

12. Rapoport, T. A., Li, L. & Park, E. Structural and Mechanistic Insights into Protein Translocation. Annu. Rev. Cell Dev. Biol. 33, 369–390 (2017).

13. Tsukazaki, T. Structural Basis of the Sec Translocon and YidC Revealed Through X-ray Crystallography. Protein J. 38, 249–261 (2019).

14. Komarudin, A. G. & Driessen, A. J. M. SecA-Mediated Protein Translocation through the SecYEG Channel. Microbiol. Spectr. 7, (2019).

15. Tsukazaki, T. et al. Structure and function of a membrane component SecDF that enhances protein export. Nature 474, 235–8 (2011).

16. Wang, P. & Dalbey, R. E. Inserting membrane proteins: the YidC/Oxa1/Alb3 machinery in bacteria, mitochondria, and chloroplasts. Biochim. Biophys. Acta 1808, 866–75 (2011).

17. Hennon, S. W., Soman, R., Zhu, L. & Dalbey, R. E. YidC/Alb3/Oxa1 Family of Insertases. J. Biol. Chem. 290, 14866–74 (2015).

18. Hegde, R. S. The Function, Structure, and Origins of the ER Membrane Protein Complex. Annu. Rev. Biochem. 91, 651–678 (2022).

19. Kumazaki, K. et al. Structural basis of Sec-independent membrane protein insertion by YidC. Nature 509, 516–20 (2014).

20. Shimokawa-Chiba, N. et al. Hydrophilic microenvironment required for the channel-independent insertase function of YidC protein. Proc. Natl. Acad. Sci. U. S. A. 112, 5063–8 (2015).

21. Sarker, S., Rudd, K. E. & Oliver, D. Revised translation start site for secM defines an atypical signal peptide that regulates Escherichia coli secA expression. J. Bacteriol. 182, 5592–5 (2000).

22. Nakatogawa, H. & Ito, K. The Ribosomal Exit Tunnel Functions as a Discriminating Gate. Cell 108, 629–636 (2002).

23. Rajapandi, T., Dolan, K. M. & Oliver, D. B. The first gene in the Escherichia coli secA operon, gene X, encodes a nonessential secretory protein. J. Bacteriol. 173, 7092–7 (1991).

24. McNicholas, P., Salavati, R. & Oliver, D. Dual regulation of Escherichia coli secA translation by distinct upstream elements. J. Mol. Biol. 265, 128–41 (1997).

25. Nakatogawa, H. & Ito, K. Secretion Monitor, SecM, Undergoes Self-Translation Arrest in the Cytosol. Mol. Cell 7, 185–192 (2001).

26. Butkus, M. E., Prundeanu, L. B. & Oliver, D. B. Translocon ‘pulling’ of nascent SecM controls the duration of its translational pause and secretion-responsive secA regulation. J. Bacteriol. 185, 6719–22 (2003).

27. Ito, K., Mori, H. & Chiba, S. Monitoring substrate enables real-time regulation of a protein localization pathway. FEMS Microbiol. Lett. 365, fny109 (2018).

28. Sakiyama, K., Shimokawa-Chiba, N., Fujiwara, K. & Chiba, S. Search for translation arrest peptides encoded upstream of genes for components of protein localization pathways. Nucleic Acids Res. 49, 1550–1566 (2021).

29. Parks, D. H. et al. A standardized bacterial taxonomy based on genome phylogeny substantially revises the tree of life. Nat. Biotechnol. 36, 996–1004 (2018).

30. Steinegger, M. & Söding, J. MMseqs2 enables sensitive protein sequence searching for the analysis of massive data sets. Nat. Biotechnol. 35, 1026–1028 (2017).

31. Shimizu, Y. et al. Cell-free translation reconstituted with purified components. Nat. Biotechnol. 19, 751–5 (2001).

32. Chiba, S. et al. Recruitment of a species-specific translational arrest module to monitor different cellular processes. Proc. Natl. Acad. Sci. U. S. A. 108, 6073–6078 (2011).

33. Hartz, D., McPheeters, D. S. & Gold, L. Selection of the initiator tRNA by Escherichia coli initiation factors. Genes Dev. 3, 1899–912 (1989).

34. Muto, H., Nakatogawa, H. & Ito, K. Genetically encoded but nonpolypeptide prolyl-tRNA functions in the A site for SecM-mediated ribosomal stall. Mol. Cell 22, 545–52 (2006).

35. Chiba, S. & Ito, K. Multisite Ribosomal Stalling: A Unique Mode of Regulatory Nascent Chain Action Revealed for MifM. Mol. Cell 47, 863–872 (2012).

36. Ismail, N., Hedman, R., Schiller, N. & von Heijne, G. A biphasic pulling force acts on transmembrane helices during translocon-mediated membrane integration. Nat. Struct. Mol. Biol. 19, 1018–22 (2012).

37. Fujiwara, K., Katagi, Y., Ito, K. & Chiba, S. Proteome-wide Capture of Co-translational Protein Dynamics in Bacillus subtilis Using TnDR, a Transposable Protein-Dynamics Reporter. Cell Rep. 33, 108250 (2020).

38. Shiota, N., Shimokawa-Chiba, N., Fujiwara, K. & Chiba, S. Identification of Bacillus subtilis YidC Substrates Using a MifM-instructed Translation Arrest-based Reporter. J. Mol. Biol. 435, 168172 (2023).

39. van der Sluis, E. O. & Driessen, A. J. M. Stepwise evolution of the Sec machinery in Proteobacteria. Trends Microbiol. 14, 105–8 (2006).

40. Debarbouille, M., Arnaud, M., Fouet, A., Klier, A. & Rapoport, G. The sacT gene regulating the sacPA operon in Bacillus subtilis shares strong homology with transcriptional antiterminators. J. Bacteriol. 172, 3966–73 (1990).

41. Arnaud, M. et al. Regulation of the sacPA operon of Bacillus subtilis: identification of phosphotransferase system components involved in SacT activity. J. Bacteriol. 174, 3161– 70 (1992).

42. Arnaud, M., Débarbouillé, M., Rapoport, G., Saier, M. H. & Reizer, J. In vitro reconstitution of transcriptional antitermination by the SacT and SacY proteins of Bacillus subtilis. J. Biol. Chem. 271, 18966–72 (1996).

43. Roy, G. et al. Posttranscriptional Regulation by Copper with a New Upstream Open Reading Frame. MBio 13, e0091222 (2022).

44. Öztürk, Y. et al. Metabolic Sensing of Extracytoplasmic Copper Availability via Translational Control by a Nascent Exported Protein. MBio 14, e0304022 (2023).

45. Tanner, D. R., Cariello, D. A., Woolstenhulme, C. J., Broadbent, M. A. & Buskirk, A. R. Genetic identification of nascent peptides that induce ribosome stalling. J. Biol. Chem. 284, 34809–18 (2009).

46. Woolstenhulme, C. J. et al. Nascent peptides that block protein synthesis in bacteria. Proc. Natl. Acad. Sci. U. S. A. 110, E878–87 (2013).

47. Sohmen, D. et al. Structure of the Bacillus subtilis 70S ribosome reveals the basis for species-specific stalling. Nat. Commun. 6, 6941 (2015).

48. Xue, L. et al. Visualizing translation dynamics at atomic detail inside a bacterial cell. Nature 610, 205–211 (2022).

49. Yap, M.-N. & Bernstein, H. D. The plasticity of a translation arrest motif yields insights into nascent polypeptide recognition inside the ribosome tunnel. Mol. Cell 34, 201–11 (2009).

50. Altschul, S. F., Gish, W., Miller, W., Myers, E. W. & Lipman, D. J. Basic local alignment search tool. J. Mol. Biol. 215, 403–10 (1990).

51. Camacho, C. et al. BLAST+: architecture and applications. BMC Bioinformatics 10, 421 (2009).

52. Teufel, F. et al. SignalP 6.0 predicts all five types of signal peptides using protein language models. Nat. Biotechnol. 40, 1023–1025 (2022).

53. Hallgren, J. et al. DeepTMHMM predicts alpha and beta transmembrane proteins using deep neural networks. bioRxiv 2022.04.08.487609 (2022) doi:10.1101/2022.04.08.487609.

54. Sonnhammer, E. L., von Heijne, G. & Krogh, A. A hidden Markov model for predicting transmembrane helices in protein sequences. Proceedings. Int. Conf. Intell. Syst. Mol. Biol. 6, 175–82 (1998).

55. Katoh, K., Misawa, K., Kuma, K. & Miyata, T. MAFFT: a novel method for rapid multiple sequence alignment based on fast Fourier transform. Nucleic Acids Res. 30, 3059– 66 (2002).

56. Katoh, K. & Standley, D. M. MAFFT multiple sequence alignment software version 7: improvements in performance and usability. Mol. Biol. Evol. 30, 772–80 (2013).

57. Paradis, E., Claude, J. & Strimmer, K. APE: Analyses of Phylogenetics and Evolution in R language. Bioinformatics 20, 289–90 (2004).

58. Letunic, I. & Bork, P. Interactive Tree Of Life (iTOL): an online tool for phylogenetic tree display and annotation. Bioinformatics 23, 127–8 (2007).

59. Gibson, D. G. et al. Enzymatic assembly of DNA molecules up to several hundred kilobases. Nat. Methods 6, 343–5 (2009).

60. Rohland, N. & Reich, D. Cost-effective, high-throughput DNA sequencing libraries for multiplexed target capture. Genome Res. 22, 939–46 (2012).

61. Mijatovic-Rustempasic, S., Frace, M. A. & Bowen, M. D. Cost-Effective Paramagnetic Bead Technique for Purification of Cycle Sequencing Products. Sequencing 2012, 1–4 (2012).

