## Supplementary Figures for "Patchy and widespread distribution of bacterial translation arrest peptides associated with the protein localization machinery"

**A**

| AP | phylum | mORF | n |
|---|---|---|---|
| SecM | Proteobacteria | *secA* | 639 |
| uA_LPPP | Planctomycetota | *secA* | 50 |
| uA_NSP-stop | Bacteroidota | *secA* | 64 |
| uA_RQH_Armatimonadales | Armatimonadota | *secA* | 3 |
| uA_RQH_Bacteroidales | Bacteroidota | *secA* | 73 |
| uA_RQH_Desulfurobacteriales | Aquificota | *secA* | 8 |
| uA_RQH_Fimbriimonadales | Armatimonadota | *secA* | 5 |
| uA_RQH_Myxococcales | Myxococcota | *secA* | 6 |
| uA_RQH_Opitutales | Verrucomicrobiota | *secA* | 3 |
| uA_RQH_Pseudomonadales | Proteobacteria | *secA* | 130 |
| uA_RQH_UBA2386 | Planctomycetota | *secA* | 3 |
| ApdA | Actinobacteriota | *secDF* | 656 |
| ApdP | Proteobacteria | *secDF* | 370 |
| VemP | Proteobacteria | *secDF* | 342 |
| uDF_DGMK-stop | Firmicutes_A | *secDF* | 2 |
| uDF_DGMK-stop | Firmicutes_B | *secDF* | 2 |
| uDF_NAP-stop | Bacteroidota | *secDF* | 403 |
| uDF_RQH_Burkholderiales | Proteobacteria | *secDF* | 3 |
| uDF_RQH_Caldicoprobacterales | Firmicutes_A | *secDF* | 5 |
| uDF_RQH_Enterobacterales | Proteobacteria | *secDF* | 44 |
| uDF_RQH_Flavobacteriales | Bacteroidota | *secDF* | 272 |
| uDF_RQH_Geobacterales | Desulfobacterota_F | *secDF* | 8 |
| uDF_RQH_Lachnospirales | Firmicutes_A | *secDF* | 4 |
| uDF_RQH_Nitrosococcales | Proteobacteria | *secDF* | 6 |
| uDF_RQH_Pseudomonadales | Proteobacteria | *secDF* | 52 |
| uDF_RQH_Steroidobacterales | Proteobacteria | *secDF* | 3 |
| uDF_RQH_UBA1845 | Planctomycetota | *secDF* | 6 |
| uDF_RQH_UBA7662 | Bacteroidota | *secDF* | 3 |
| ApcA | Actinobacteriota | *yidC* | 1599 |
| MifM | Firmicutes | *yidC* | 592 |
| uC_KYxIW | Firmicutes_A | *yidC* | 27 |

**B**

| AP | phylum | mORF | n |
|---|---|---|---|
| uA_RQH_Acidobacteriales | Acidobacteriota | *secA* | 1 |
| uA_RQH_Desulfovibrionales | Desulfobacterota | *secA* | 1 |
| uA_RQH_Treponematales | Spirochaetota | *secA* | 1 |
| uA_RQH_Verrucomicrobiales | Verrucomicrobiota | *secA* | 1 |
| uA_RQH_Vicinamibacterales | Acidobacteriota | *secA* | 2 |
| uDF_RQH_B143-G9 | Planctomycetota | *secDF* | 1 |
| uDF_RQH_Campylobacterales | Campylobacterota | *secDF* | 1 |
| uDF_RQH_Chitinophagales | Bacteroidota | *secDF* | 2 |
| uDF_RQH_Chthoniobacterales | Verrucomicrobiota | *secDF* | 1 |
| uDF_RQH_Desulfuromonadales | Desulfobacterota_F | *secDF* | 1 |
| uDF_RQH_SBR1031 | Chloroflexota | *secDF* | 1 |
| uDF_RQH_UBA2392 | Planctomycetota | *secDF* | 2 |
| uDF_RQH_UBA8108 | Planctomycetota | *secDF* | 2 |
| uDF_RQH_WSZJ01 | Planctomycetota | *secDF* | 2 |

**Supplementary Figure 1: General information of candidate monitoring substrates**.

**(A)** Names of previously and currently identified arrest peptides (AP), their phylogenetic distributions (phylum), and their downstream genes (mORF) are listed. 'n' represents the number of ORFs identified in this study. **(B)** Orphan uORFs with RAPP/RGPP motif that essentially met our screening criteria except for their conservation among species within an individual bacterial order (n<3).

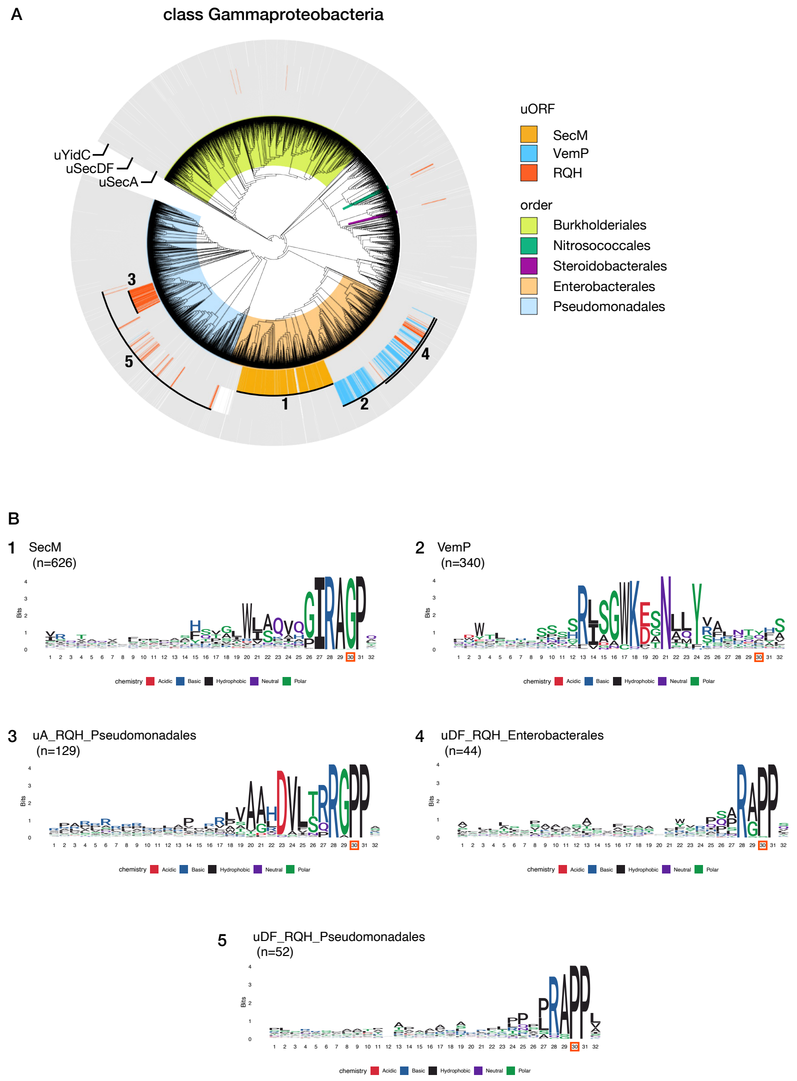


**Supplementary Figure 2: Phylogenetic distribution of candidate monitoring substrates in Gammaproteobacteria.**

**(A)** Phylogenetic distribution of the candidate monitoring substrates in class Gammaproteobacteria. The bacterial genomes that encode SecA, SecD/F, or YidC homolog are indicated by grey strips around the bacterial phylogenetic tree. Genomes that possess genes for known monitoring substrates and putative arrest peptides upstream of *secA*, *secDF*, or *yidC* gene are represented by chromatic strips with names or numbering corresponding to those shown in (B). The colors behind the tree indicate the bacterial orders listed on the right. **(B)** Sequence logos of known and candidate monitoring substrates. P-site residues at the 30$^{th}$ position (red square) with its N-terminal 29 residues, as well as the following two C-terminal residues, are used to plot sequence logos.

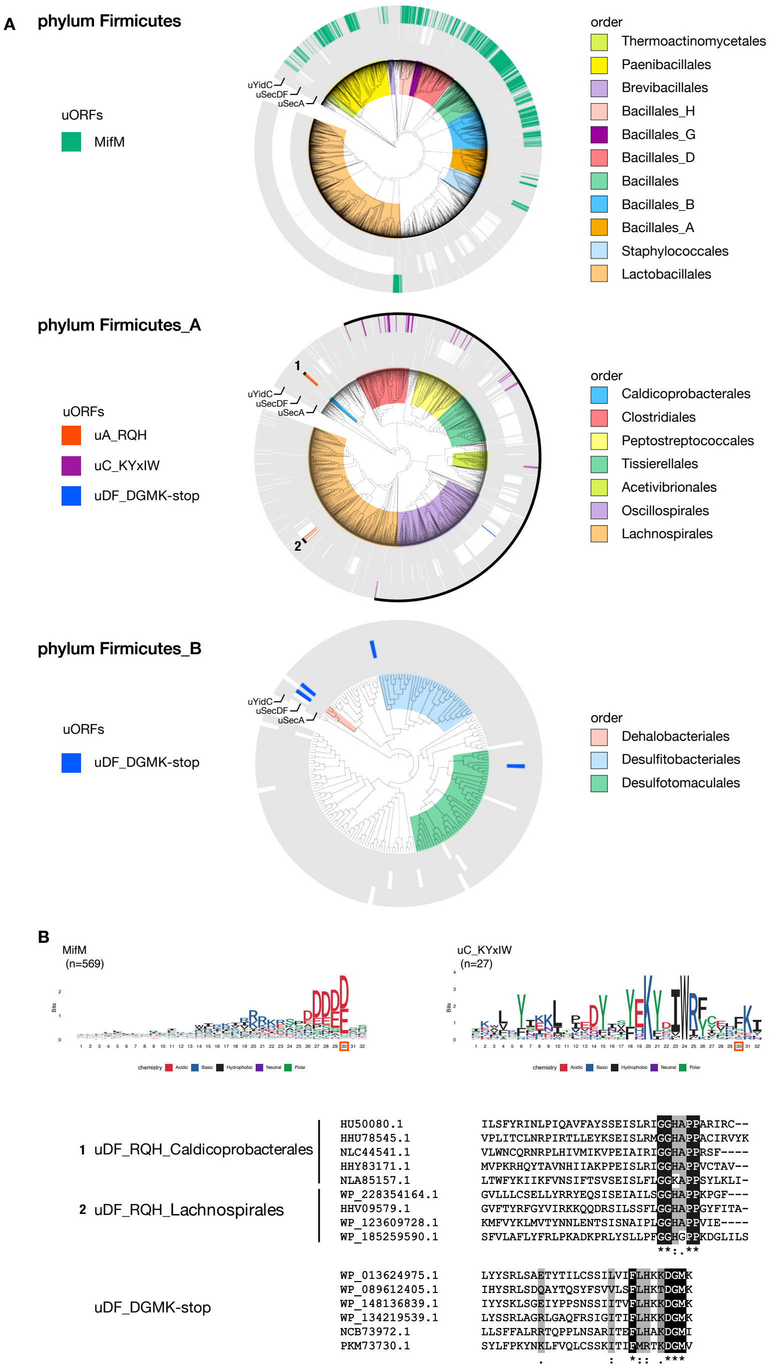


**Supplementary Figure 3: Phylogenetic distribution of candidate monitoring substrates in phylum Firmicutes.**

**(A)** Phylogenetic distribution of the known and candidate monitoring substrates in the phylum Firmicutes, Firmicutes_A and Firmicutes_B. **(B)** Sequence logos and alignments of known and candidate monitoring substrates in the phylum Firmicutes, Firmicutes_A, and Firmicutes_B. The figures follow the format used in Supplementary Fig. 2.

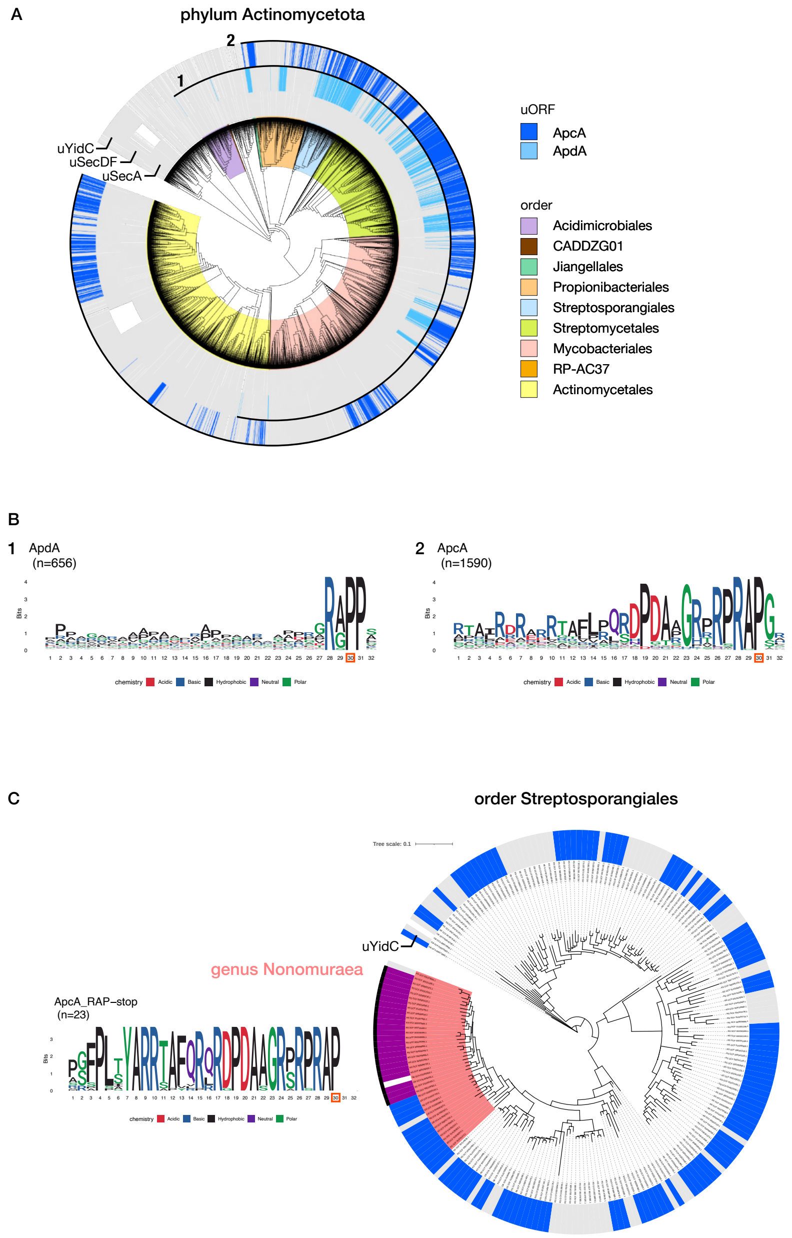


**Supplementary Figure 4: Phylogenetic distribution of candidate monitoring substrates in phylum Actinomycetota.**

**(A)** Phylogenetic distribution of the candidate monitoring substrates in phylum Actinomycetota. **(B)** Sequence logos of ApdA and ApcA. **(C)** Phylogenetic tree of order Streptosporangiales and a sequence logo of ApcA with RAP-stop found among genus Nonomuraea.

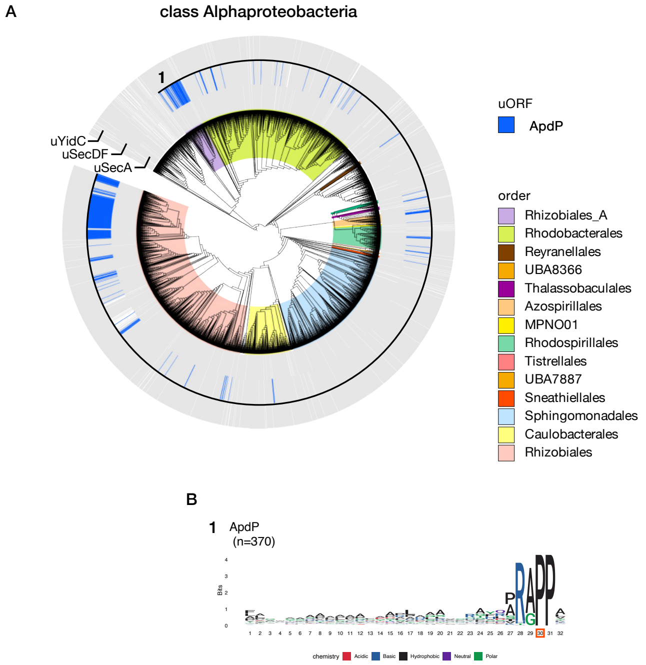


**Supplementary Figure 5: Phylogenetic distribution of ApdP in class Alphaproteobacteria. (A, B)** Phylogenetic distribution (A) and sequence logo (B) of ApdP. The figures follow the format used in Supplementary Fig. 2.

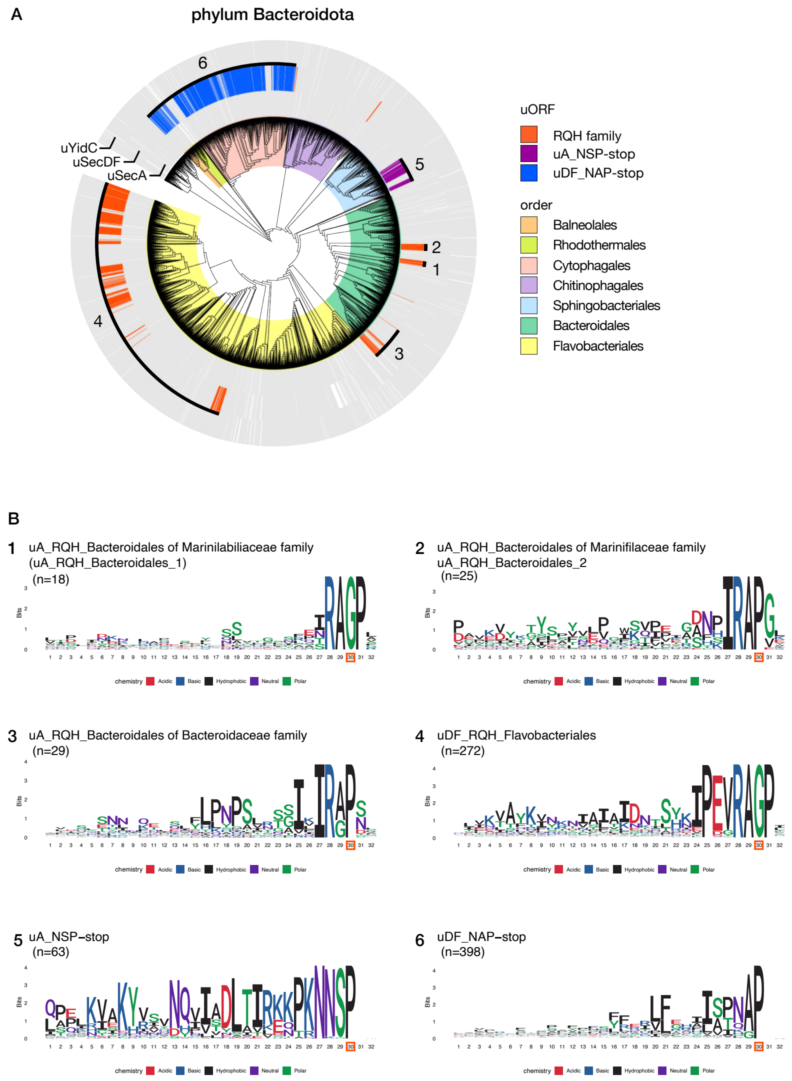


**Supplementary Figure 6: Phylogenetic distribution of candidate monitoring substrates in phylum Bacteroidota.**

**(A)** Phylogenetic distribution of the candidate monitoring substrates in phylum Bacteroidota.

**(B)** Sequence logos of candidate monitoring substrates. The figures follow the format used in Supplementary Fig. 2.

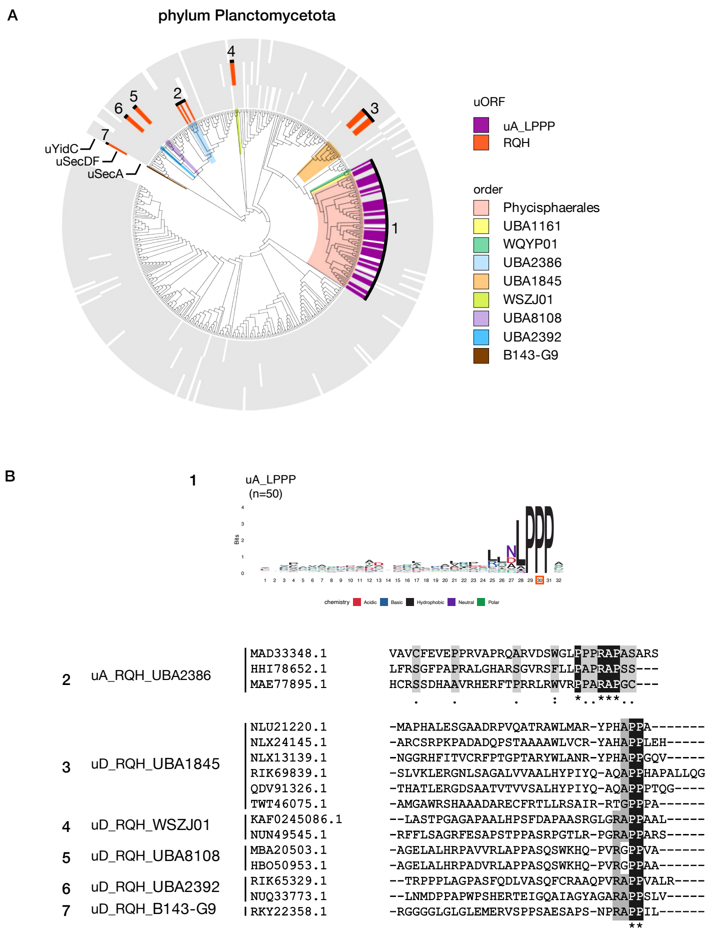


**Supplementary Figure 7: Phylogenetic distribution of candidate monitoring substrates in phylum Planctomycetota.**

**(A)** Phylogenetic distribution of the candidate monitoring substrates in phylum Planctomycetota. **(B)** A sequence logo and sequence alignments of candidate monitoring substrates. The figures follow the format used in Supplementary Fig. 2.

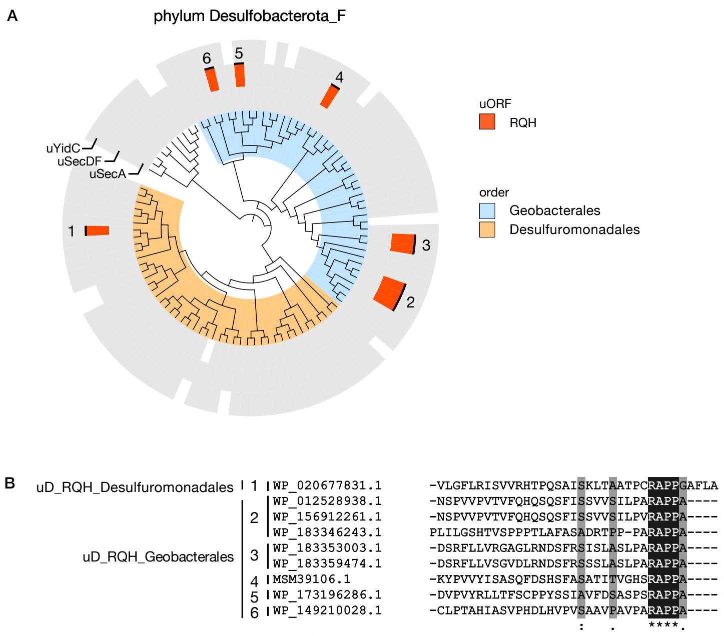


**Supplementary Figure 8: Phylogenetic distribution of candidate monitoring substrates in phylum Desulfobacterota_F.**

**(A)** Phylogenetic distribution of the candidate monitoring substrates in phylum Desulfobacterota_F. **(B)** Sequence alignment of candidate monitoring substrates. The figures follow the format used in Supplementary Fig. 2.

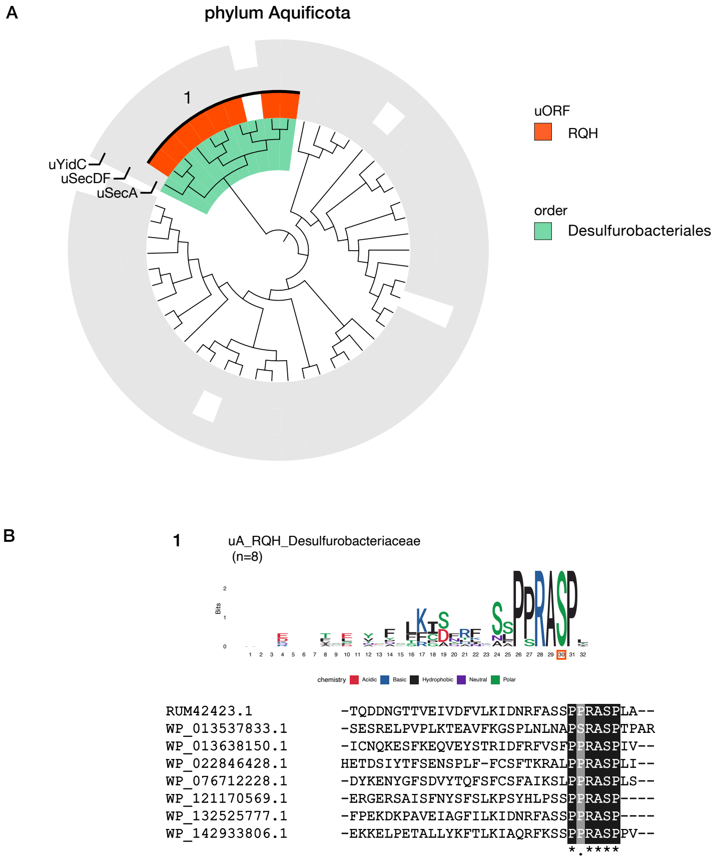


**Supplementary Figure 9: Phylogenetic distribution of candidate monitoring substrates in phylum Aquificota.**

**(A)** Phylogenetic distribution of the candidate monitoring substrates in phylum Aquificota. **(B)** A sequence logo and alignment of candidate monitoring substrates. The figures follow the format used in Supplementary Fig. 2.

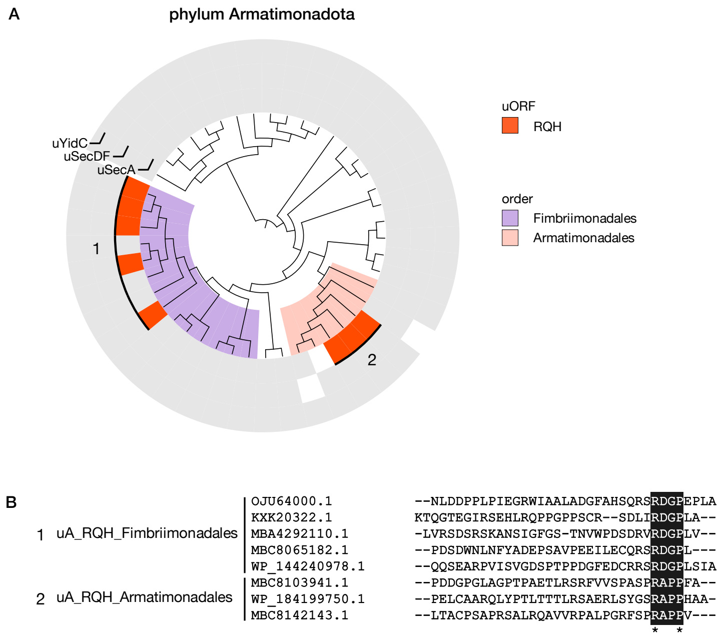


**Supplementary Figure 10: Phylogenetic distribution of candidate monitoring substrates in phylum Armatimonadota.**

(A) Phylogenetic distribution of the candidate monitoring substrates in phylum Armatimonadota. (B) A sequence alignment of candidate monitoring substrates. The figures follow the format used in Supplementary Fig. 2.

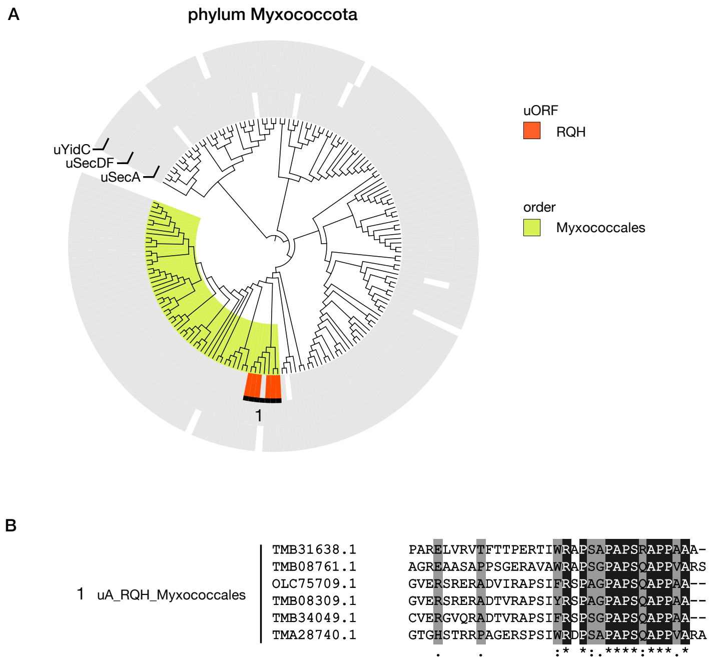


**Supplementary Figure 11: Phylogenetic distribution of candidate monitoring substrates in phylum Myxococcota.**

**(A)** Phylogenetic distribution of the candidate monitoring substrates in phylum Myxococcota. **(B)** A sequence alignment of candidate monitoring substrates. The figures follow the format used in Supplementary Fig. 2.

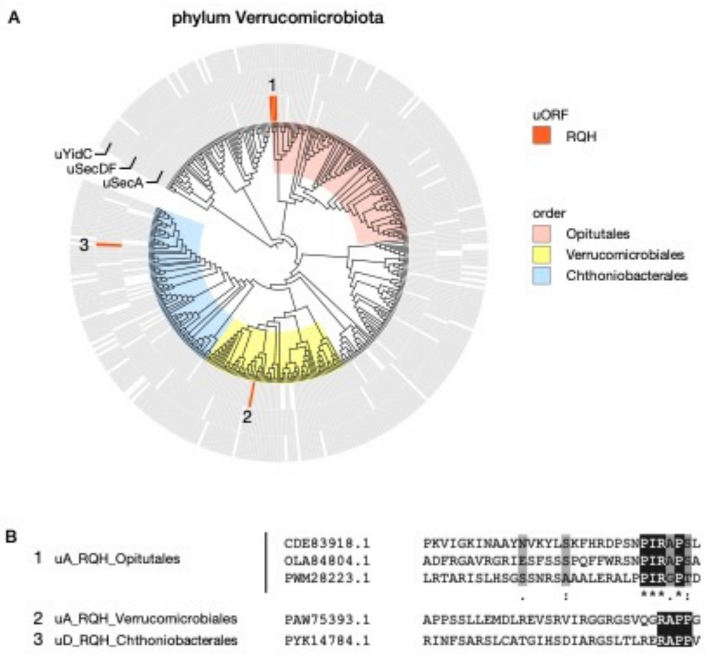


**Supplementary Figure 12: Phylogenetic distribution of candidate monitoring substrates in phylum Verrucomicrobiota.**

**(A)** Phylogenetic distribution of the candidate monitoring substrates in phylum Verrucomicrobiota. **(B)** Sequence alignment of candidate monitoring substrates. The figures follow the format used in Supplementary Fig. 2.

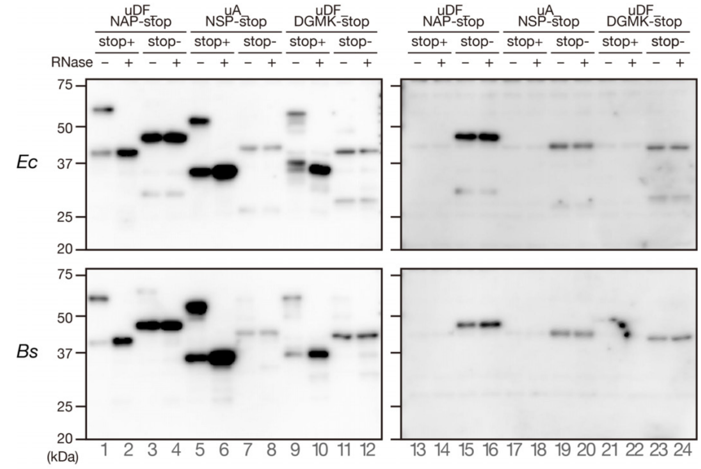


**Supplementary Figure 13: The translation arrest of uDF_NAP-stop, uA_NSP-stop and uDF_DGMK-stop depends on the stop codon.**

The *gfp-ap-lacZa* reporters harboring uDF_NAP-stop, uA_NSP-stop and uDF_DGMK-stop C-terminal region with (stop+) or without (stop-) stop codon just after the NAP, NSP or DGMK motif were translated in *Ec* (upper) or *Bs* (lower) PURE. The translation products were immunoblotted using anti-GFP (left membranes) or anti-LacZα (right membranes). Before electrophoresis, portions of the sample were treated by RNase A (lanes indicated as +) to degrade tRNA moiety.

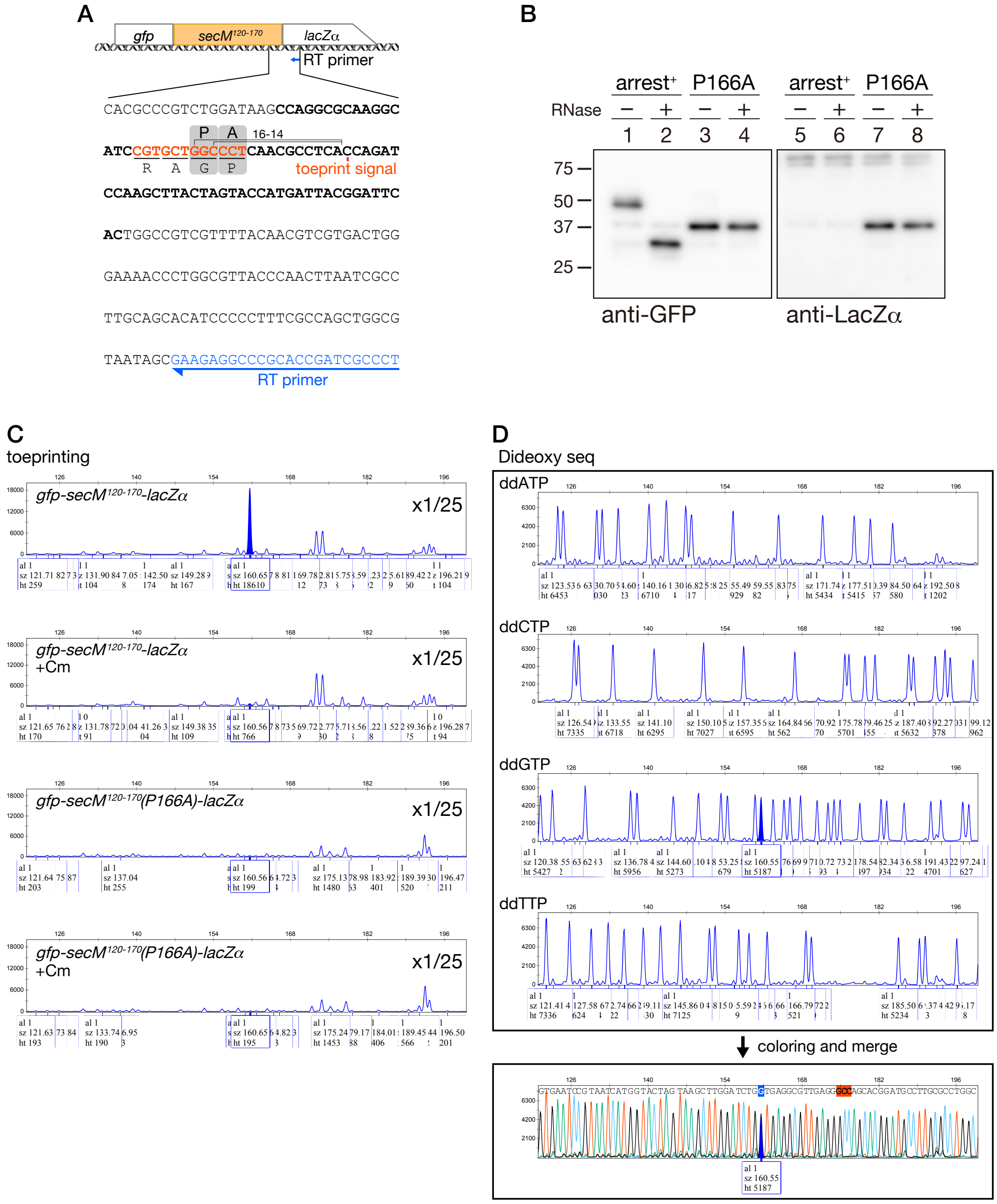


**Supplementary Figure 14: Identification of the ribosome stalling site of *E. coli* SecM by toeprinting.**

**(A)** A schematic representation of the template gene for in vitro translation, in which the coding region for SecM residues 120-170 was sandwich-fused between *gfp* and *lacZα* (upper). The sequence between the coding region for SecM arrest site and the downstream primer annealing site for reverse transcription and dideoxy sequencing is shown at the bottom. The residue corresponding to the toeprint signal (results shown in C and D) and the estimated ribosome stalling site are shown. The gray boxes with 'P' and 'A' indicate P- and A-site codons, respectively. **(B)** Western blotting analysis of translation product of the reporter by *Ec* PURE system. Before electrophoresis, portions of the sample were treated by RNase A (lanes indicated as +) to degrade tRNA moiety. **(C)** Raw results of the fragment analysis of toeprint product using a capillary sequencer. The template gene shown in (A) was translated by *E*c PURE with or without chloramphenicol, and subsequently subjected to reverse transcription using the reverse transcription (RT) primer shown in (A). The size of the complementary DNA was then analyzed by the fragment analysis using a capillary DNA sequencer. The stalling-dependent toeprint signal specifically detected from the experiment using the wild-type but not from the arrest-defective P166A derivative of SecM in the absence of chloramphenicol is filled in blue. **(D)** Raw results of the fragment analysis of dideoxy sequencing products using the same RT primer. Each plot was colored (A; green, C; right blue, G; black, T; red) and then overlayed as shown at the bottom. The peaks whose fragment size are same as that of the stalling-dependent toeprint signal were filled in blue. The toeprint product generated by the SecM-stalled ribosome appeared as a single peak in our fragment analysis with the size indicating that the ribosome stalled with the P-site at the $Gly_{165}$ codon and the A-site at the $Pro_{166}$ codon (Fig. 3B and S14).

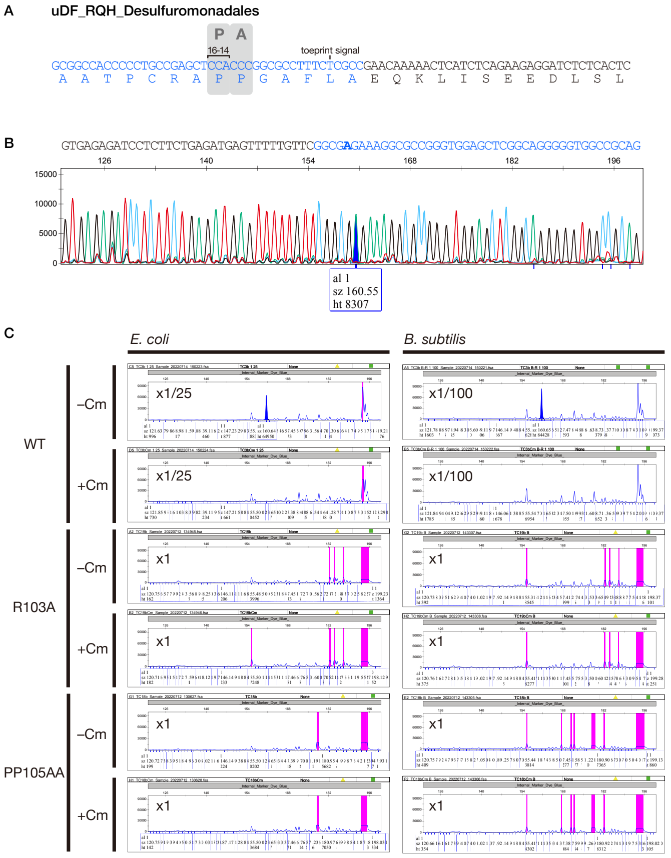


**Supplementary Figure 15: Identification of the ribosome stalling site of uDF_RQH_Desulfuromonadales by toeprinting.**

**(A)** A schematic representation of the ribosome stalling site of uDF_RQH_Desulfuromonadales estimated by toeprinting. The nucleotide and amino acid sequences derived from the coding region of uDF_RQH_Desulfuromonadales are shown in blue characters. The estimated ribosome stalling site was shown by the P-site (P) and A-site (A) codons. **(B)** Overlayed results of dideoxy sequencing. The reverse complement sequence of the *uDF_RQH_Desulfuromonadales* region is presented in blue, and the peak corresponding to the stalling-specific toeprint signal and its nucleotide in the dideoxy sequencing data are shown by the filled blue peak and the bold alphabet, respectively. The numbers above the peak data indicate the estimated sizes (nucleotides) based on a molecular weight standard (500LIZ). **(C)** Raw results of the fragment analysis of toeprint products yielded by the reverse transcription after in vitro translation of wild-type (WT) and arrest-defective mutant derivatives (R103A, PP105AA) of uDF_RQH_Desulfuromonadales using *Ec* (left) and *Bs* (right) PURE. Each template DNA was subjected to translation in the presence (+Cm) or absence (-Cm) of chloramphenicol. When necessary, the toeprinting products were diluted with HiDi formamide prior to the capillary electrophoresis at the ratios indicated in each panel. The magenta lines represent saturated peaks.

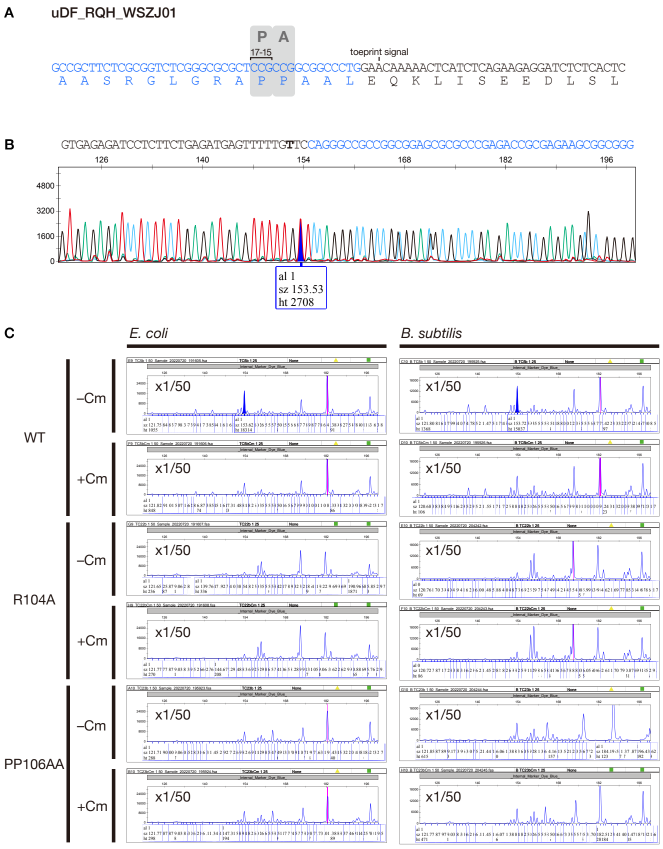


**Supplementary Figure 16: Identification of the ribosome stalling site of uDF_RQH_WSZJ01 by toeprinting.**

**(A)** A schematic representation of the ribosome stalling site of uDF_RQH_WSZJ01 estimated by toeprinting. **(B)** Overlayed results of dideoxy sequencing. The peak corresponding to the stalling-specific toeprint signal and its nucleotide in the dideoxy sequencing data are shown by the filled blue peak and the bold alphabet, respectively. **(C)** Raw results of toeprinting analysis of wild-type and arrest-defective mutant derivatives. Detailed information is described in the legend of Supplementary Fig. 15.

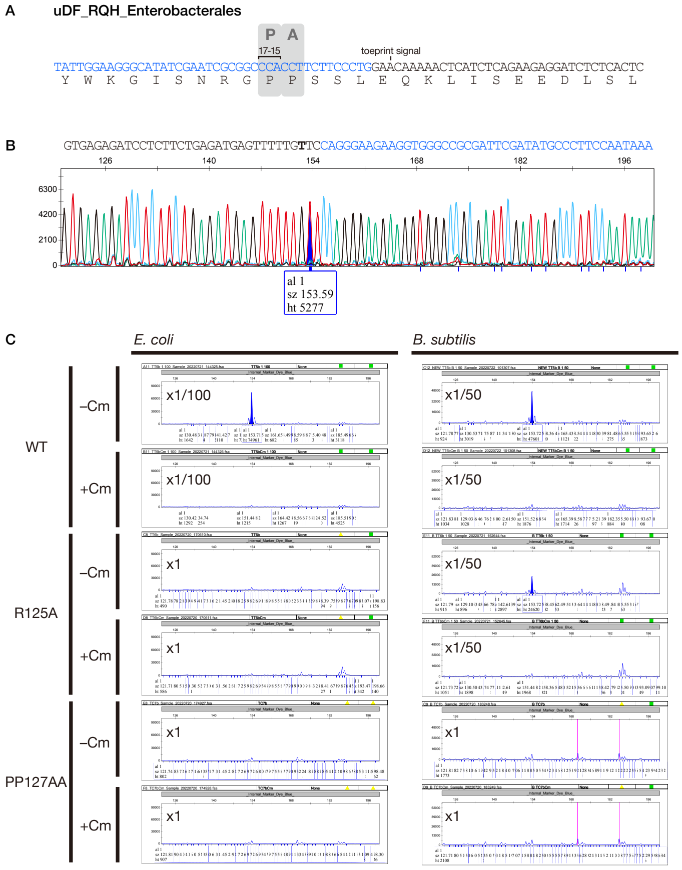


**Supplementary Figure 17: Identification of the ribosome stalling site of uDF_RQH_Enterobacterales by toeprinting.**

**(A)** A schematic representation of the ribosome stalling site of uDF_RQH_Enterobacterales estimated by toeprinting. **(B)** Overlayed results of dideoxy sequencing. The peak corresponding to the stalling-specific toeprint signal and its nucleotide in the dideoxy sequencing data are shown by the filled blue peak and the bold alphabet, respectively. **(C)** Raw results of toeprinting analysis of wild-type and arrest-defective mutant derivatives. Detailed information is described in the legend of Supplementary Fig. 15.

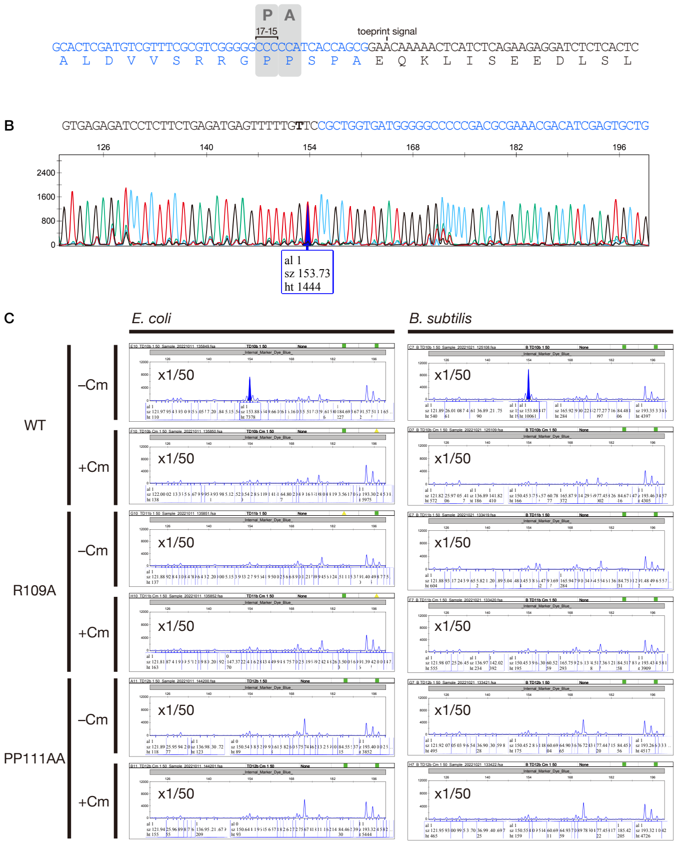


**Supplementary Figure 18: Identification of the ribosome stalling site of uA_RQH_Pseudomonadales by toeprinting.**

**(A)** A schematic representation of the ribosome stalling site of uA_RQH_Pseudomonadales estimated by toeprinting. **(B)** Overlayed results of dideoxy sequencing. The peak corresponding to the stalling-specific toeprint signal and its nucleotide in the dideoxy sequencing data are shown by the filled blue peak and the bold alphabet, respectively. **(C)** Raw results of toeprinting analysis of wild-type and arrest-defective mutant derivatives. Detailed information is described in the legend of Supplementary Fig. 15.

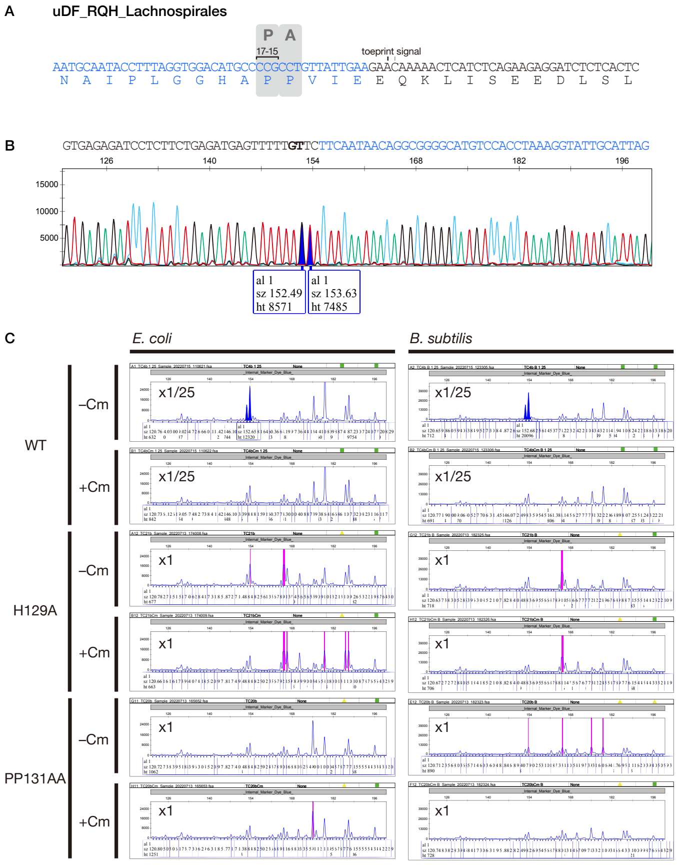


**Supplementary Figure 19: Identification of the ribosome stalling site of uDF_RQH_Lachnospirales by toeprinting.**

**(A)** A schematic representation of the ribosome stalling site of uDF_RQH_Lachnospirales estimated by toeprinting. **(B)** Overlayed results of dideoxy sequencing. The peaks corresponding to the stalling-specific toeprint signals and their nucleotides in the dideoxy sequencing data are shown by the filled blue peak and the bold alphabets, respectively. **(C)** Raw results of toeprinting analysis of wild-type and arrest-defective mutant derivatives. Detailed information is described in the legend of Supplementary Fig. 15.

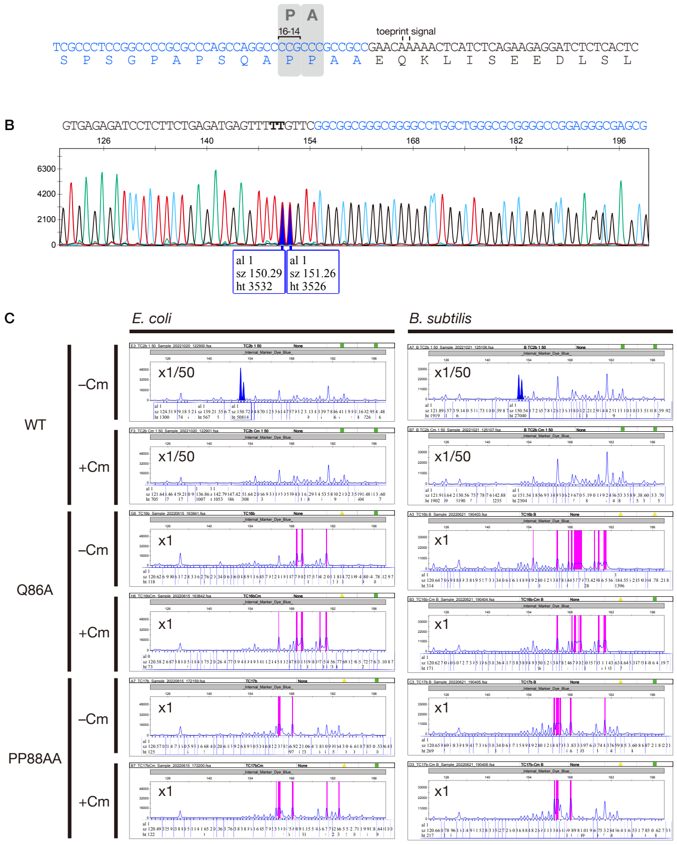


**Supplementary Figure 20: Identification of the ribosome stalling site of uA_RQH_Myxococcales by toeprinting.**

**(A)** A schematic representation of the ribosome stalling site of uA_RQH_Myxococcales estimated by toeprinting. **(B)** Overlayed results of dideoxy sequencing. The peaks corresponding to the stalling-specific toeprint signals and their nucleotides in the dideoxy sequencing data are shown by the filled blue peak and the bold alphabet, respectively. **(C)** Raw results of toeprinting analysis of wild-type and arrest-defective mutant derivatives. Detailed information is described in the legend of Supplementary Fig. 15.

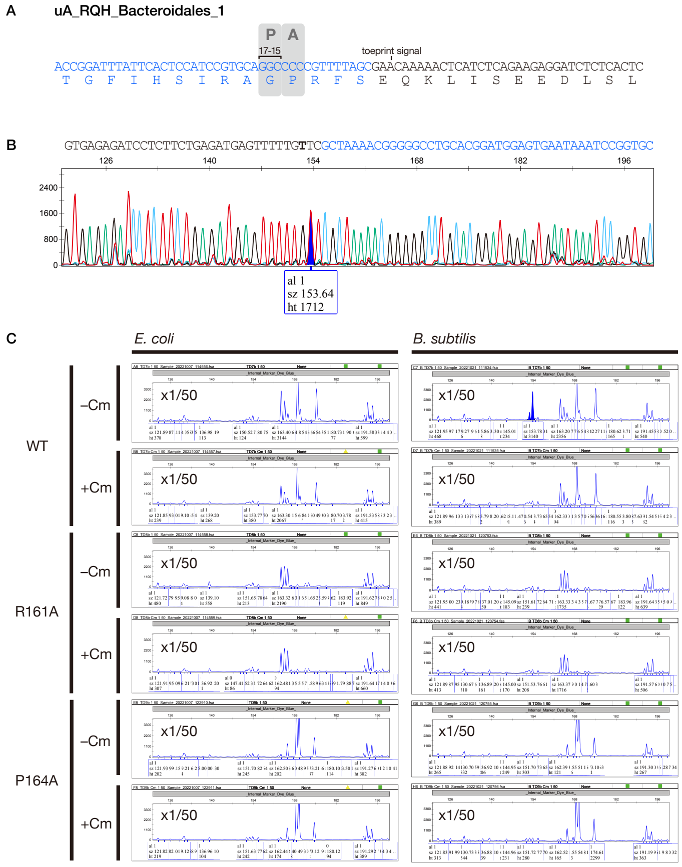


**Supplementary Figure 21: Identification of the ribosome stalling site of uA_RQH_Bacteroidales_1 by toeprinting.**

**(A)** A schematic representation of the ribosome stalling site of uA_RQH_Bacteroidales_1 estimated by toeprinting. **(B)** Overlayed results of dideoxy sequencing. The peak corresponding to the stalling-specific toeprint signal and its nucleotide in the dideoxy sequencing data are shown by the filled blue peak and the bold alphabet, respectively. **(C)** Raw results of toeprinting analysis of wild-type and arrest-defective mutant derivatives. Detailed information is described in the legend of Supplementary Fig. 15.

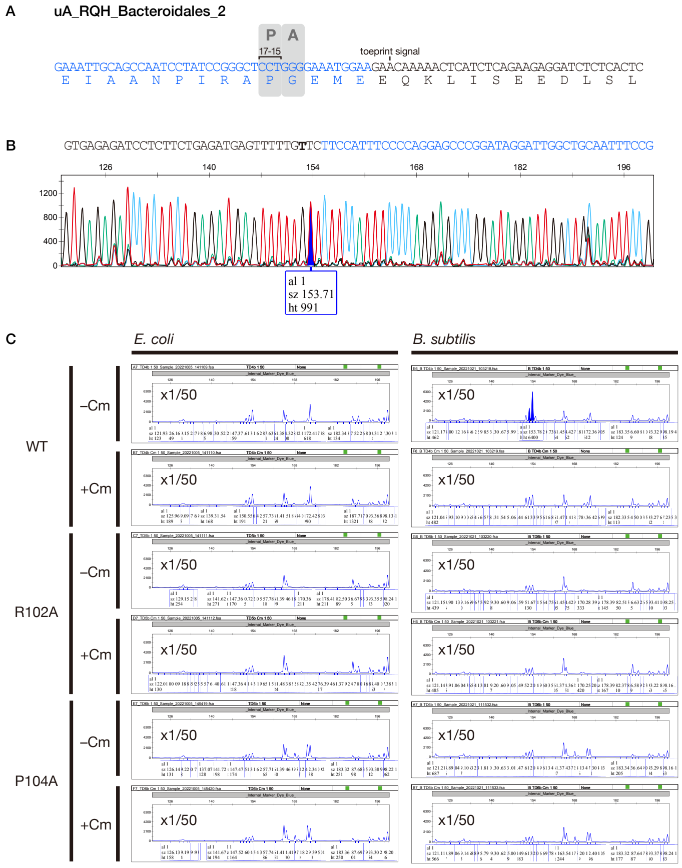


**Supplementary Figure 22: Identification of the ribosome stalling site of uA_RQH_Bacteroidales_2 by toeprinting.**

**(A)** A schematic representation of the ribosome stalling site of uA_RQH_Bacteroidales_2 estimated by toeprinting. **(B)** Overlayed results of dideoxy sequencing. The peak corresponding to the stalling-specific toeprint signal and its nucleotide in the dideoxy sequencing data are shown by the filled blue peak and the bold alphabet, respectively. **(C)** Raw results of toeprinting analysis of wild-type and arrest-defective mutant derivatives. Detailed information is described in the legend of Supplementary Fig. 15.

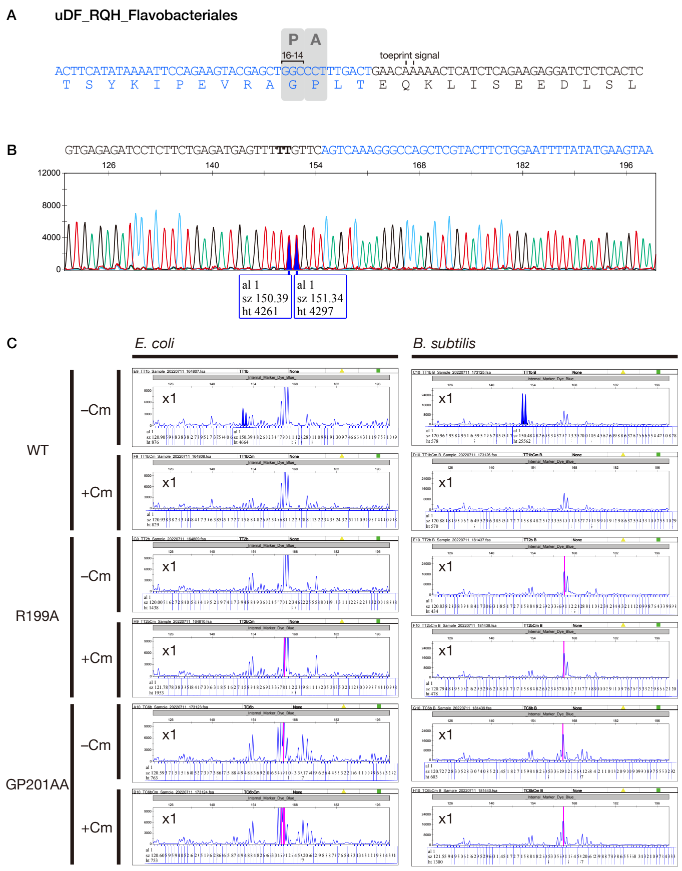


**Supplementary Figure 23: Identification of the ribosome stalling site of uDF_RQH_Flavobacteriales by toeprinting.**

**(A)** A schematic representation of the ribosome stalling site of uDF_RQH_Flavobacteriales estimated by toeprinting. **(B)** Overlayed results of dideoxy sequencing. The peaks corresponding to the stalling-specific toeprint signals and their nucleotides in the dideoxy sequencing data are shown by the filled blue peak and the bold alphabet, respectively. **(C)** Raw results of toeprinting analysis of wild-type and arrest-defective mutant derivatives. Detailed information is described in the legend of Supplementary Fig. 15.

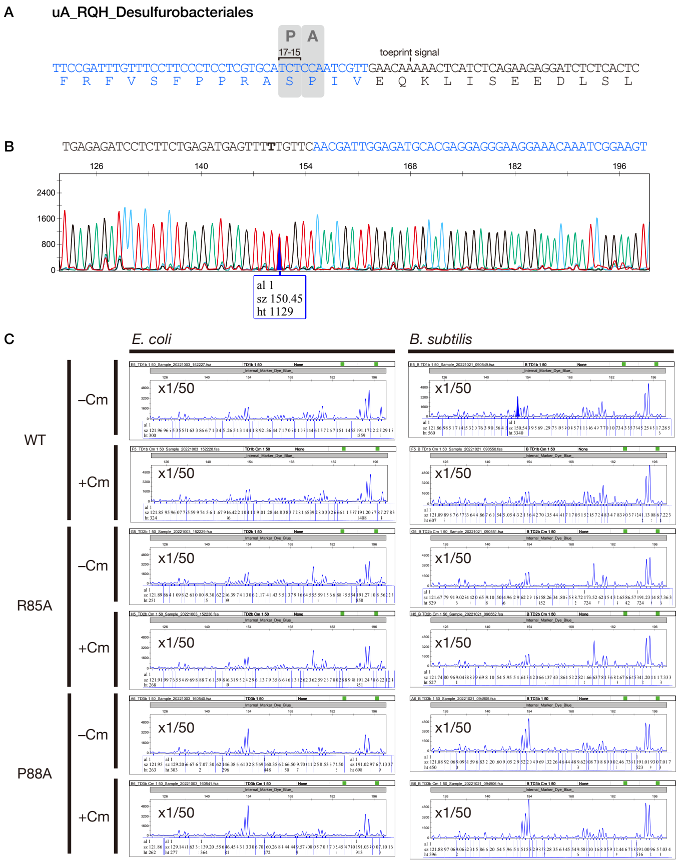


**Supplementary Figure 24: Identification of the ribosome stalling site of uA_RQH_Desulfurobacteriales by toeprinting.**

**(A)** A schematic representation of the ribosome stalling site of uA_RQH_Desulfurobacteriales estimated by toeprinting. **(B)** Overlayed results of dideoxy sequencing. The peak corresponding to the stalling-specific toeprint signal and its nucleotide in the dideoxy sequencing data are shown by the filled blue peak and the bold alphabet, respectively. **(C)** Raw results of toeprinting analysis of wild-type and arrest-defective mutant derivatives. Detailed information is described in the legend of Supplementary Fig. 15.

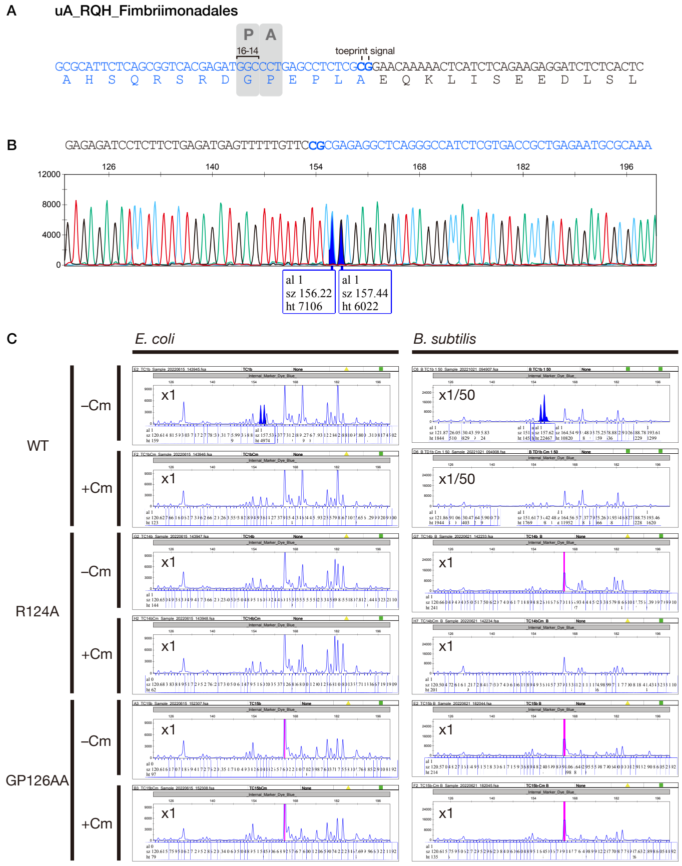


**Supplementary Figure 25: Identification of the ribosome stalling site of uA_RQH_Fimbriimonadales by toeprinting.**

**(A)** A schematic representation of the ribosome stalling site of uA_RQH_Fimbriimonadales estimated by toeprinting. **(B)** Overlayed results of dideoxy sequencing. The peaks corresponding to the stalling-specific toeprint signals and their nucleotides in the dideoxy sequencing data are shown by the filled blue peak and the bold alphabet, respectively. **(C)** Raw results of toeprinting analysis of wild-type and arrest-defective mutant derivatives. Detailed information is described in the legend of Supplementary Fig. 15.

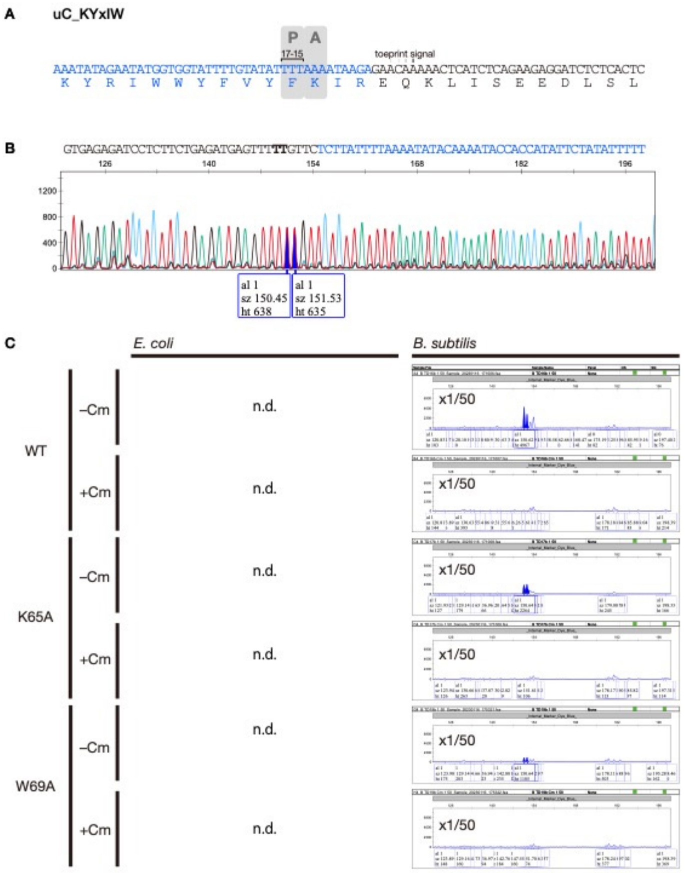


**Supplementary Figure 26: Identification of the ribosome stalling site of uC_KYxIW by toeprinting.**

**(A)** A schematic representation of the ribosome stalling site of uC_KYxIW estimated by toeprinting. **(B)** Overlayed results of dideoxy sequencing. The peaks corresponding to the stalling-specific toeprint signals and their nucleotides in the dideoxy sequencing data are shown by the filled blue peak and the bold alphabet, respectively. **(C)** Raw results of toeprinting analysis of wild-type and arrest-defective mutant derivatives. Detailed information is described in the legend of Supplementary Fig. 15.

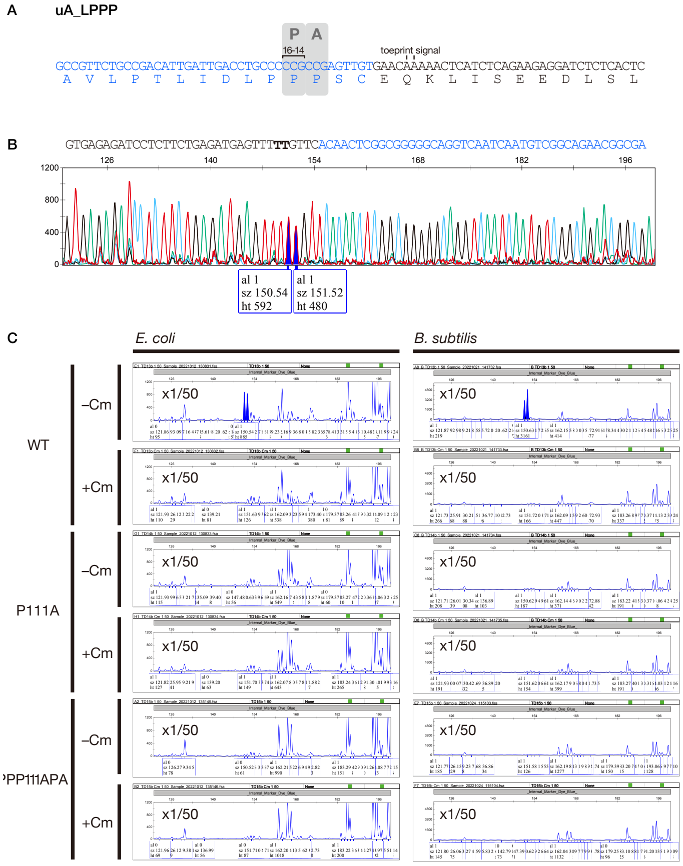


**Supplementary Figure 27: Identification of the ribosome stalling site of uA_LPPP by toeprinting.**

**(A)** A schematic representation of the ribosome stalling site of uA_LPPP estimated by toeprinting. **(B)** Overlayed results of dideoxy sequencing. The peaks corresponding to the stalling-specific toeprint signals and their nucleotides in the dideoxy sequencing data are shown by the filled blue peak and the bold alphabet, respectively. **(C)** Raw results of toeprinting analysis of wild-type and arrest-defective mutant derivatives. Detailed information is described in the legend of Supplementary Fig. 15.

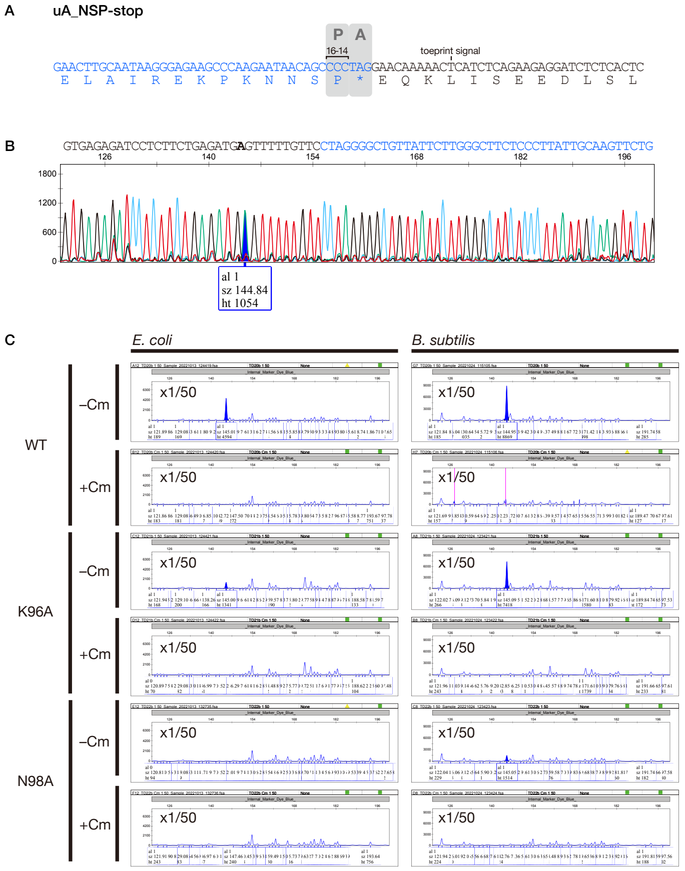


**Supplementary Figure 28: Identification of the ribosome stalling site of uA_NSP-stop by toeprinting.**

**(A)** A schematic representation of the ribosome stalling site of uA_NSP-stop estimated by toeprinting. **(B)** Overlayed results of dideoxy sequencing. The peak corresponding to the stalling-specific toeprint signal and its nucleotide in the dideoxy sequencing data are shown by the filled blue peak and the bold alphabet, respectively. **(C)** Raw results of toeprinting analysis of wild-type and arrest-defective mutant derivatives. Detailed information is described in the legend of Supplementary Fig. 15.

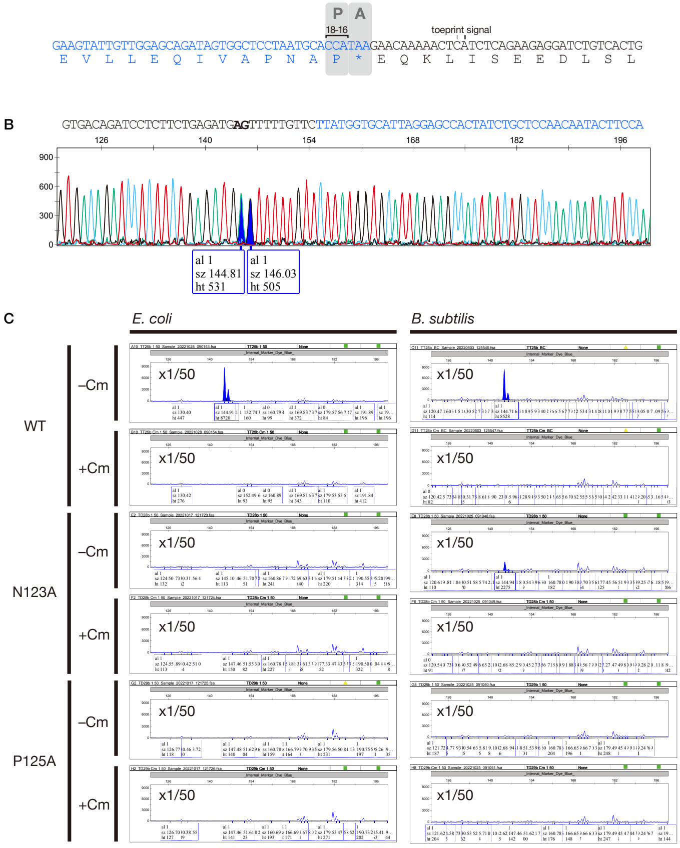


**Supplementary Figure 29: Identification of the ribosome stalling site of uDF_NAP-stop by toeprinting.**

**(A)** A schematic representation of the ribosome stalling site of uDF_NAP-stop estimated by toeprinting. **(B)** Overlayed results of dideoxy sequencing. The peaks corresponding to the stalling-specific toeprint signals and their nucleotides in the dideoxy sequencing data are shown by the filled blue peak and the bold alphabet, respectively. **(C)** Raw results of toeprinting analysis of wild-type and arrest-defective mutant derivatives. Detailed information is described in the legend of Supplementary Fig. 15.

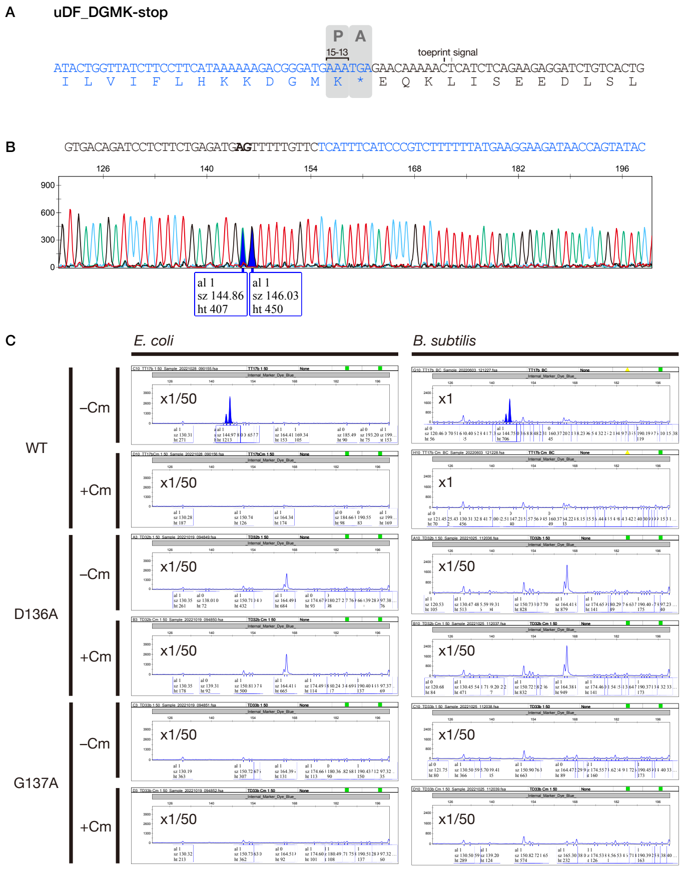


**Supplementary Figure 30: Identification of the ribosome stalling site of uDF_DGMK-stop by toeprinting.**

**(A)** A schematic representation of the ribosome stalling site of uDF_DGMK-stop estimated by toeprinting. **(B)** Overlayed results of dideoxy sequencing. The peaks corresponding to the stalling-specific toeprint signals and their nucleotides in the dideoxy sequencing data are shown by the filled blue peak and the bold alphabet, respectively. **(C)** Raw results of toeprinting analysis of wild-type and arrest-defective mutant derivatives. Detailed information is described in the legend of Supplementary Fig. 15.

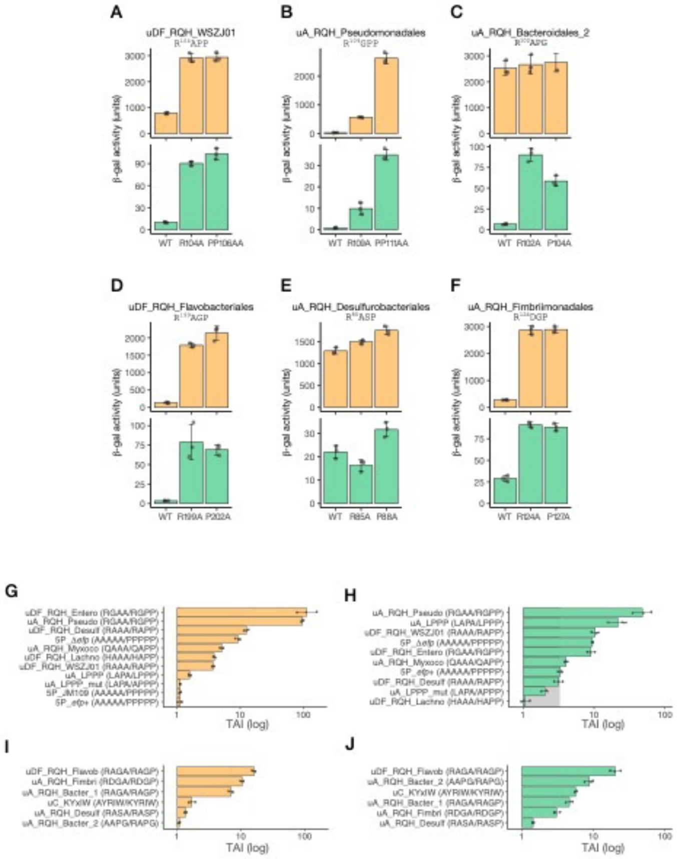


**Supplementary Figure 31: In vivo translation arrest assay of candidate monitoring substrates**

**(A-F)** the β-galactosidase activity of *E. coli* (orange bars) and *B. subtilis* cells (green bars), harboring wild-type (WT) or mutant derivatives of arrest peptide reporters. **(G-J)** Translation arrest indexes (TAI) are calculated based on the in vivo β-galactosidase activities of mutant reporter divided by those of wild-type reporter. Error bars (mean ± standard deviations, n = 3) and individual data points (dots) are indicated.

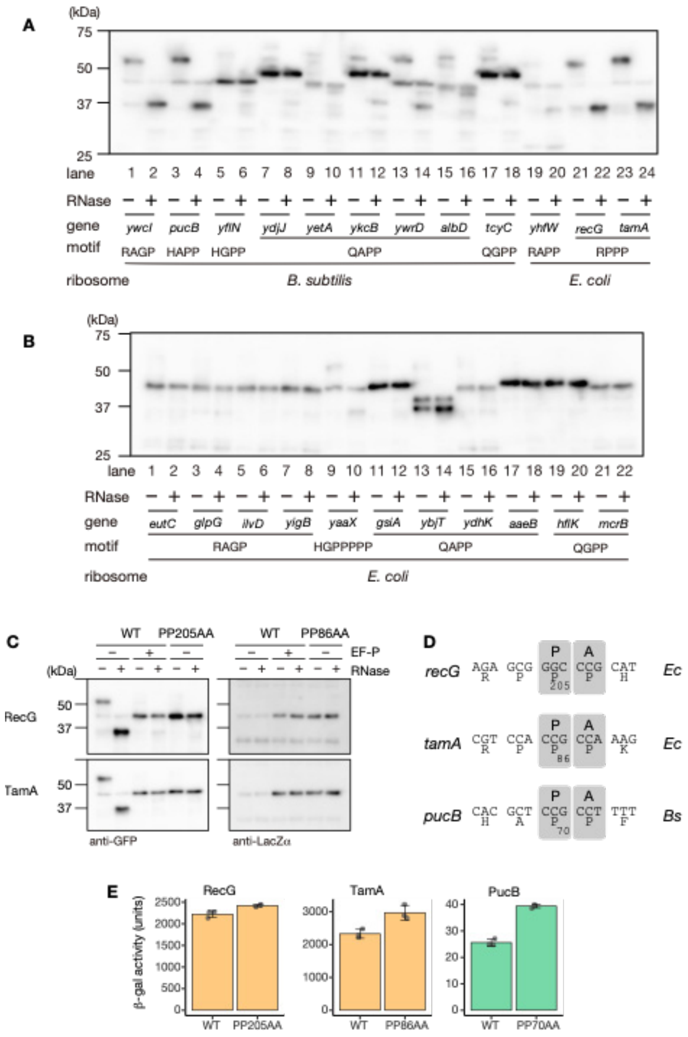

**Supplementary Figure 32: Investigation for translation arrest by amino acid sequences harboring RAPP-like motifs derived from *E. coli* and *B. subtilis* proteome.** The genes tested are listed in **Supplementary** Table 5. **(A-C)** Western blotting of in vitro translation products. The *gfp-ap-lacZα* reporters were translated in *Ec* or *Bs* PURE. The translation products were then separated by neutral pH gels and immunoblotted using anti-GFP or anti-LacZα. Before the separation, portions of samples were treated by RNase A (lanes indicated as +) to degrade tRNA moiety. **(D)** Estimated ribosome stalling sites based on the results obtained by toeprinting analysis. **(E)** In vivo translation arrest assay. β-galactosidase activity of *E. coli* (orange bars) and *B. subtilis* cells (green bars), harboring wild-type (WT) or mutant derivatives of arrest peptide reporters. Error bars (mean ± standard deviations, n = 3) and individual data points (dots) are indicated.

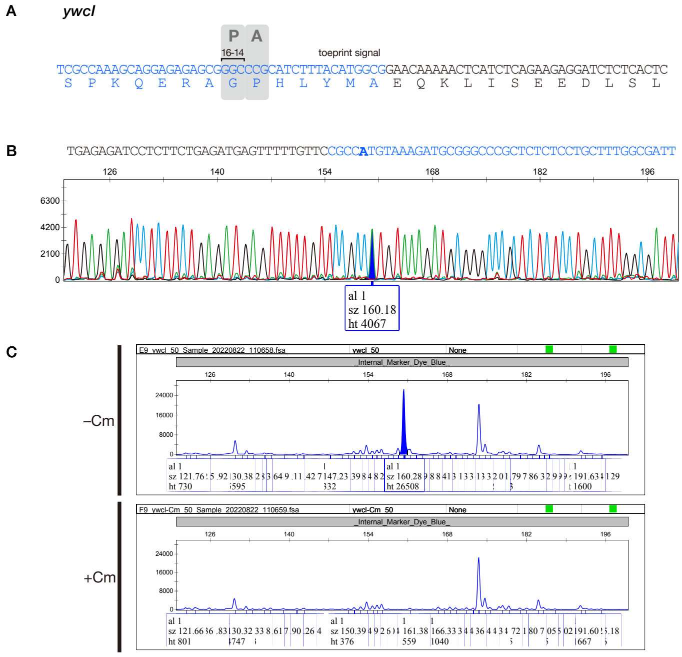


**Supplementary Figure 33: Identification of the ribosome stalling site of *ywcI* by toeprinting.**

**(A)** A schematic representation of the ribosome stalling site of *ywcI* estimated by toeprinting. **(B)** Overlayed results of dideoxy sequencing. The peak corresponding to the stalling-specific toeprint signal and its nucleotide in the dideoxy sequencing data are shown by the filled blue peak and the bold alphabet, respectively. **(C)** Raw results of toeprinting analysis of wild-type and arrest-defective mutant derivatives. Detailed information is described in the legend of Supplementary Fig. 15.
